# ComBat Harmonization: Empirical Bayes versus Fully Bayes Approaches

**DOI:** 10.1101/2022.07.13.499561

**Authors:** Maxwell Reynolds, Tigmanshu Chaudhary, Mahbaneh Eshaghzadeh Torbati, Dana L. Tudorascu, Kayhan Batmanghelich, the Alzheimer’s Disease Neuroimaging Initiative

## Abstract

*Studying small effects or subtle neuroanatomical variation requires large-scale sample size data. As a result, combining neuroimaging data from multiple datasets is necessary. Variation in acquisition protocols, magnetic field strength, scanner build, and many other non-biologically related factors can introduce undesirable bias into studies. Hence, harmonization is required to remove the bias-inducing factors from the data. ComBat is one of the most common methods applied to features from structural images. ComBat models the data using a hierarchical Bayesian model and uses the empirical Bayes approach to infer the distribution of the unknown factors. The empirical Bayes harmonization method is computationally efficient and provides valid point estimates. However, it tends to underestimate uncertainty. This paper investigates a new approach, fully Bayesian ComBat, where Monte Carlo sampling is used for statistical inference. When comparing fully Bayesian and empirical Bayesian ComBat, we found Empirical Bayesian ComBat more effectively removed scanner strength information and was much more computationally efficient. Conversely, fully Bayesian ComBat better preserved biological disease and age-related information while performing more accurate harmonization on traveling subjects. The fully Bayesian approach generates a rich posterior distribution, which is useful for generating simulated imaging features for improving classifier performance in a limited data setting. We show the generative capacity of our model for augmenting and improving the detection of patients with Alzheimer’s disease. Posterior distributions for harmonized imaging measures can also be used for brain-wide uncertainty comparison and more principled downstream statistical analysis. Code for our new fully Bayesian ComBat extension is available at https://github.com/batmanlab/BayesComBat*.

## 1. Introduction

Large-scale neuroimaging datasets have been created in recent years to identify disease biomarkers, study brain development, and standardize image acquisition (Mueller et al., 2005). These datasets have enabled the identification of disease-relevant features (King et al., 2009), population-wide examination of neurological phenotypes (Cury et al., 2015), individual brain trajectory modeling (Koval et al., 2021), and data-driven disease subtyping (Young et al., 2018). The open-access nature of these datasets has allowed for external validation of new findings (Cury et al., 2020), and large longitudinal datasets have enabled subject-specific prediction with ground truth validation (Marinescu et al., 2020; Nebli et al., 2020).

Projects like the Alzheimer’s Disease Neuroimaging Initiative (ADNI) (Mueller et al., 2005) and the Adolescence Brain Cognitive Development (ABCD) Study (Casey et al., 2018) have led to insights into Alzheimer’s Disease, aging, development, and other biological processes. Most of these datasets use images acquired from many different clinical sites and scanners. Differences in magnetic resonance imaging (MRI) scanner hardware and acquisition processes from these multiple sites can introduce additional unwanted variance into neuroanatomical feature measurements (Han et al., 2006). For example, a 3T scanner may produce a higher quality tissue contrast than a 1.5T scanner, leading to higher estimates of grey matter volumes. Quantifying and correcting for nonbiological scanner factors, while maintaining biological information, is necessary to facilitate more accurate analysis so that scanner effects are not attributed to subject or population differences (Fortin et al., 2017). Harmonization addresses this issue by modeling and correcting for scanner effects in imaging features.

Many methods have been proposed for harmonizing raw images and imaging-derived features (e.g. regional grey matter thickness and volume). For image-level harmonization, recent work has viewed harmonization as a style transfer procedure and used deep learning approaches including variational auto-encoders (Zuo et al., 2021), generative adversarial networks (GANs) (Liu et al., 2021), and U-Nets (Torbati et al., 2021b) to harmonize images to a specific reference scanner. Deep learning image-based approaches are increasingly adopted for various applications in the medical domain, including harmonization and subsequent downstream tasks such as disease classification. However, feature-based methods that measure regional and global brain characteristics such as thickness and volume remain relevant for studies of neurological disease pathology and progression due to their interpretable nature. Such features are readily related to the biological understanding of disease models. We therefore focus on image-derived feature harmonization and build on an existing widely used harmonization method.

Feature-level harmonization is often treated as a regression problem. One approach is to model scanner effects for each imaging feature as a fixed effect and residualize the effect from the data (Venkatraman et al., 2015). Another method, ME-Mega, uses a similar model but views scanner effects as random intercepts (Radua et al., 2020). ComBat, a batch harmonization technique originally proposed for gene expression microarrays (Johnson et al., 2007), adds a multiplicative (variance scaling) scanner effect term. Additionally, ComBat assumes that site effects come from a common distribution across regions of interest by using a hierarchical Bayesian model. This causes pooling of scanner effects towards a mean, making ComBat more robust to smaller within-scanner sample sizes (Johnson et al., 2007).

ComBat has recently been proposed for structural MRI-derived feature harmonization (Fortin et al., 2018, 2017) and has since been used routinely for harmonization in neuroimaging studies (Bartlett et al., 2018; Dima et al., 2021; Habes et al., 2021). The original ComBat model has also been further extended to accommodate repeated scans on the same subjects over time in longitudinal datasets such as ADNI (Beer et al., 2020).

ComBat uses a type of Bayesian inference called empirical Bayes (EB) (Carlin and Louis, 2000) to infer the distribution of the latent variables. In EB, the observed data is used to learn a point estimation of latent variables at the highest level of the hierarchical model (hyperparameters), rather than learning a probability distribution. Empirical Bayes is often computationally less expensive, especially for large models such as ComBat, but the hyperparameter point estimation ignores uncertainty in part of the model. This can lead to inaccurate and underestimated posterior uncertainty for the latent variables (van de Wiel et al., 2019). Additionally, the empirical Bayes approach confines the posterior of a parameter of interest to a specific distribution (e.g., Normal or Inverse-Gamma). For models with conjugate priors, this assumption is valid. However, this limits the choice of a prior distribution. Using a fully Bayesian approach generally produces more accurate uncertainty measurements (Gelman et al., 2021; van de Wiel et al., 2019), allowing for more accurate posterior distribution inference even when some model parameters are misspecified (Piecuch et al., 2017).

Using fully Bayesian approaches has typically relied on slower Markov Chain Monte Carlo (MCMC) inference methods such as Metropolis-Hastings MCMC (Hastings, 1970) for inference of the posterior distribution of the model’s latent variables. The sampling approach allows more flexible choice of prior distributions at the expense of the computational cost of inference. Recently however, more efficient samplers and parallel GPU computation have enabled computationally feasible fully Bayesian estimation for large models (Phan et al., 2019). As the efficiency difference between empirical and fully Bayesian inference narrows, fully Bayesian inference may offer a more principled approach without prohibitive computational costs.

With this work, we contribute to the literature in multiple ways: 1) We introduce a new ComBat formulation which infers a joint posterior distribution for the entire model in a single inference stage; 2) we investigate the performance for harmonization of features from T1-weighted structural images against EB ComBat using metrics to quantify biological (e.g. age and disease) information while removing non-biological information (e.g. scanner strength and test-retest feature differences); and 3) we introduce several novel use cases for FB harmonization which utilize its rich posterior distribution for augmentation and uncertainty quantification.

## 2. Materials and Methods

### 2.1 Data

We use T1-weighted structural images from the ADNI dataset, acquired using MPRAGE on Philips, Siemens, and GE scanners (Jack et al., 2010). ADNI is a public-private partnership launched in 2003 with the primary goal of testing if combinations of MRI, positron emission tomography (PET), other biomarkers, and clinical and neuropsychological assessments can measure MCI and AD progression. More information can be found at www.adni-info.org (ADNI, 2016). All subjects gave informed consent in accordance with local Institutional Review Board Regulations (Petersen et al., 2010).

We obtain 3894 initial images from 809 patients grouped as either cognitively normal (CN), mild cognitive impaired (MCI), or Alzheimer’s disease (AD). At baseline, 28.1% (n=227) of subjects are labeled CN, 49.1% (n=397) are labeled MCI, and 22.9% (n=185) are labeled AD. The subjects’ sexes are 57% (n=467) male and 43.2% (n=342) female. The subjects’ ages at baseline are between 51 and 91 years with a mean age of 75.3 years. Images were acquired on 83 different scanners. 58 of the scanners have a 1.5T field strength; the remaining 25 scanners are 3T.

### 2.2 Preprocessing

We use the FreeSurfer version 7.1.1 longitudinal pipeline (Reuter et al., 2012) on a Linux CentOS version 8.2 machine to segment various brain structures and obtain global and local cortical thickness and subcortical volume measurements. The first step in the Freesurfer Longitudinal pipeline is the standard cross-sectional “recon-all” function. This includes motion correction, N3 non-uniformity correction, brain extraction, subcortical segmentation, and cortical parcellation of each image. Next, a mean template from each within-subject image set is created and used for an unbiased initialization for a second “recon-all” run on each image.

### 2.3 Quality Control

After FreeSurfer processing, 401 images are dropped due to duplicate scans (subject scanned on the same scanner on the same date) or failure during FreeSurfer registration, segmentation, or parcellation stages. Next, 70 images with outlier imaging features (at least 5 standard deviations away from the feature mean) are manually inspected and excluded if noticeable errors existed such as brain extraction failure leading to segmentations labeling the skull as cortical gray matter.

### 2.4 Empirical Bayes ComBat Model

The EB ComBat model (Beer et al., 2020) is given by:

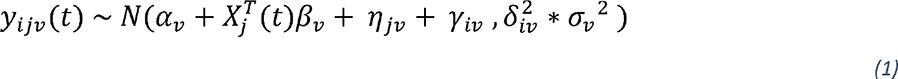

where *i* is the scanner index, *j* is the subject index, *v* is the imaging feature index, and *t* represents time. *j*_*ijv*_(*t*) is the measured (unharmonized) value for feature *v* of subject *i* on scanner *j*. *γ*_*iv*_ is the additive scanner factor for scanner *i* and feature *v*. 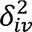 is the scaling scanner factor from scanner *i* and feature *v*. 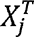 is a covariate term for biological effects. A description of all variables is given in Table 1. We include age, sex, diagnosis, and diagnosis x age covariate terms in both EB and FB ComBat models (Beer et al., 2020; Sun et al., 2021). Of note, the subject-specific random effect *η*_*jv*_ was added in longitudinal ComBat (Beer et al., 2020) and showed improved harmonization in longitudinal datasets. We therefore include this parameter in both our EB and FB models.

**Table 1:**
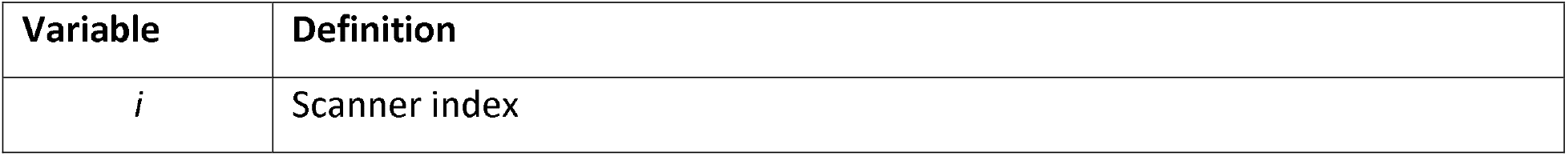
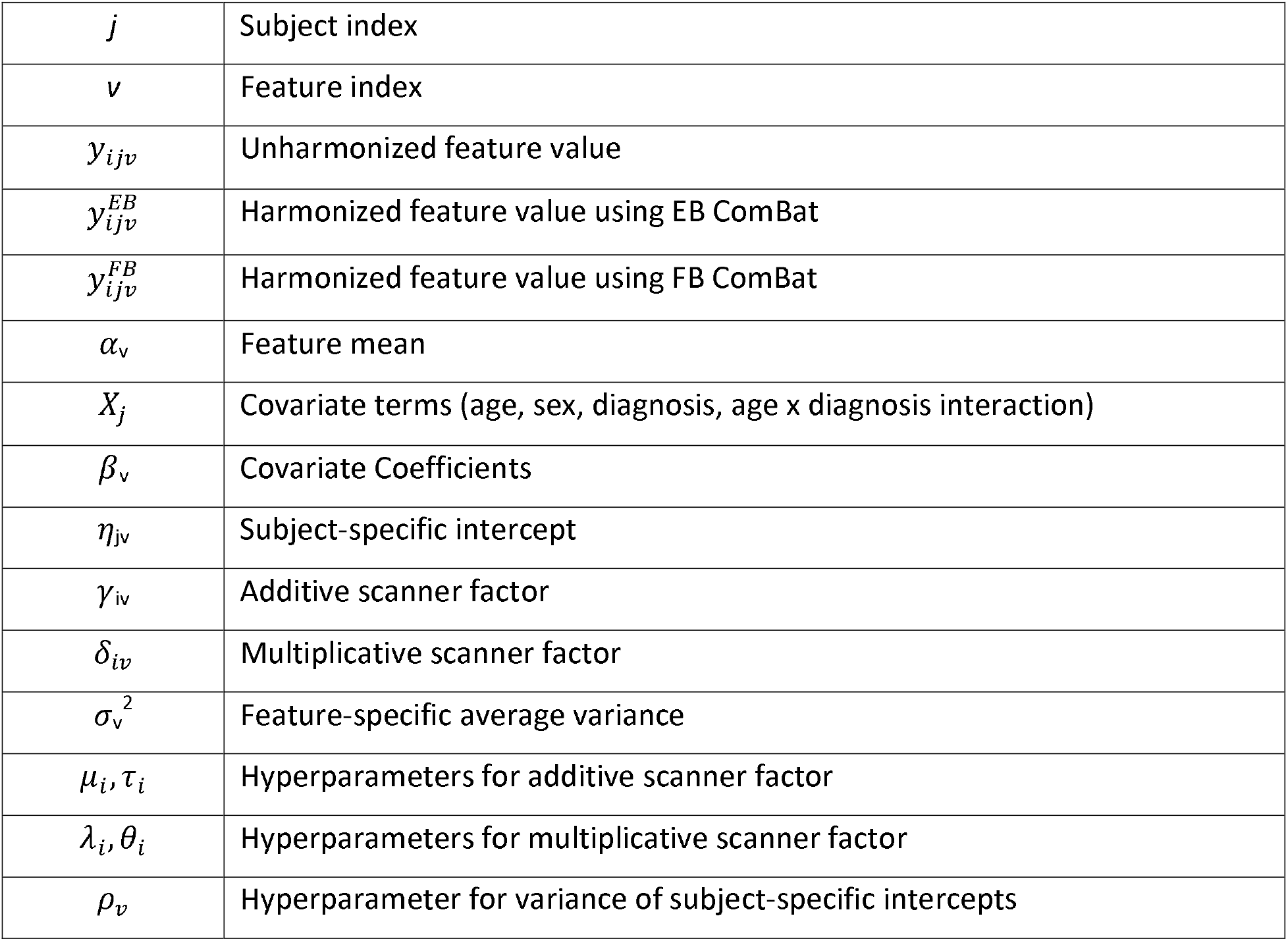
ComBat equation variables

In EB ComBat, *α_v_, β_v_, η_jv_, σ_v_*, and priors for *γ_iv_* and 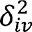 are estimated using restricted maximum likelihood (REML) and method of moments, then conditional posteriors for *γ_iv_* and 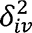 are identified using an expectation-maximization (EM) algorithm.

Harmonized adjusted feature values follow the equation:

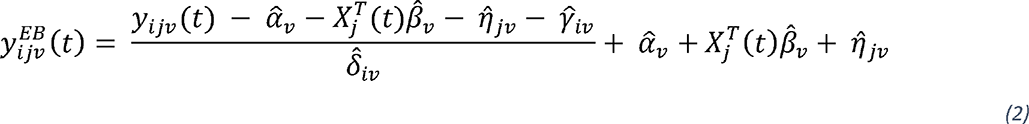

where 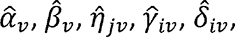 are the parameter estimates.

### 2.5 Fully Bayes ComBat Model

In FB ComBat, all high-level parameters are given weakly informative hyper-priors. A plate diagram including hyperparameters is shown in Figure 1. We choose prior distributions to be centered at 0 for additive factors and 1 for variance parameters. Fat-tailed Cauchy and Half-Cauchy distributions are used for several parameters when values close to 0 are expected with the possibility of outliers (e.g. a very biased scanner additive factor).

**Figure 1:**
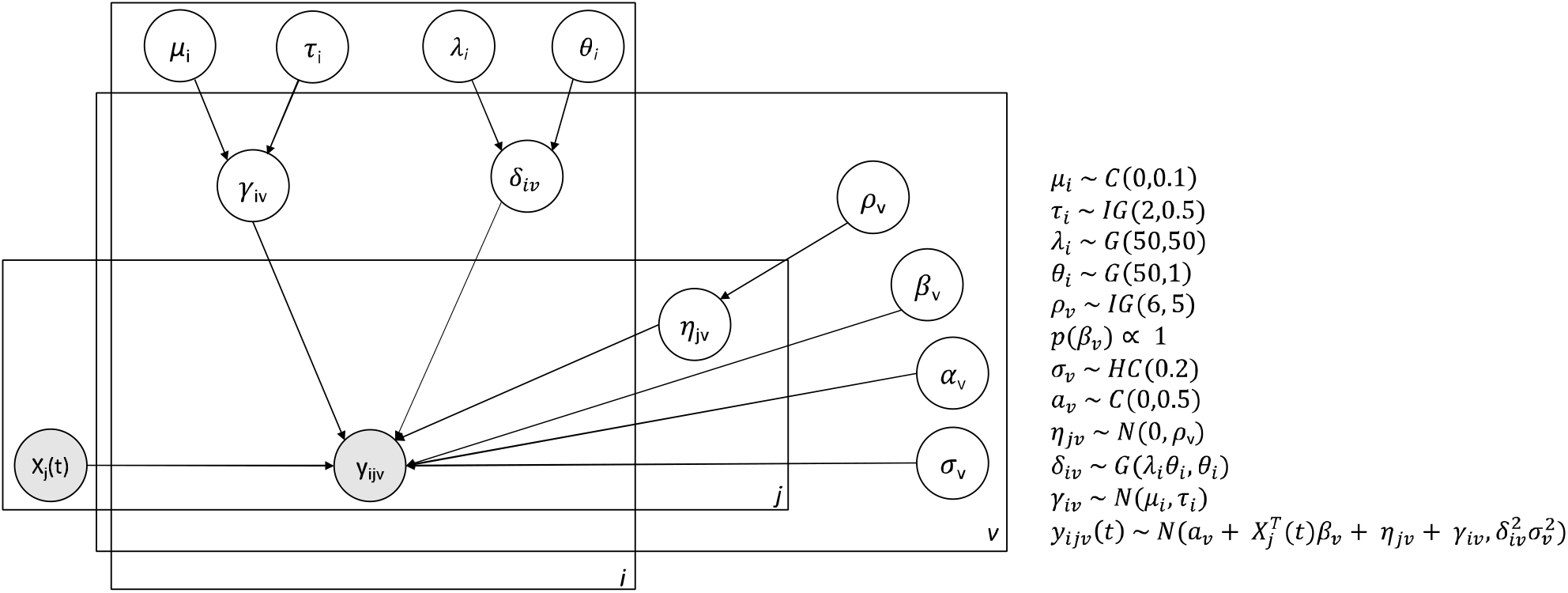
FB ComBat Plate Diagram Plate diagram for FB ComBat model. Shaded circles represent observed measurements (covariates and imaging feature values). Unshaded circles represent latent parameters. Distributions C, IG, G, HC, N are Cauchy, Inverse Gamma, Gamma, Half-Cauchy, and Normal respectively.

We add the following constraints to scanner additive and multiplicative factors to ensure identifiability:

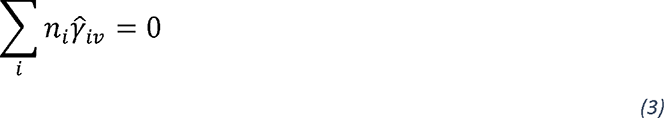

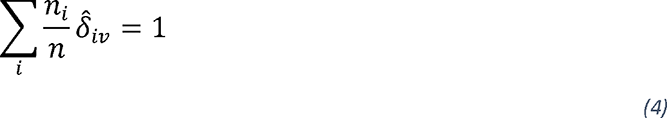

where *n* is the total number of images, and *n_i_* is the number of images from scanner *i*. The identifiability constraint on additive scanner parameters *(3)* is explicitly made in EB ComBat. The constraint on multiplicative scanner parameters *(4)* ensures that the average overall error variance of imaging features is not changed by harmonization. In EB ComBat, this constraint is made implicitly by a multi-step inference procedure of first estimating *σ_v_*, then learning *δ_iv_* and rescaling the features accordingly (Beer et al., 2020). FB ComBat samples all parameters jointly, so in this approach, we perform a transformation on the *σ_v_* and *δ_iv_* samples which preserves overall error variance while ensuring *(4)* is met.

Features are standardized to have a mean of 0 and a standard deviation of 1 before inference. Hamiltonian Monte Carlo (HMC) using a No-U-Turn Sampler (NUTS) (Hoffman and Gelman, 2014) is performed for 40,000 samples on the entire model, yielding posterior distribution samples for all model parameters. Briefly, HMC sampling models the negative log model posterior space as a *p*-dimensional physical space where *p* is the number of parameters in a model. Sampling trajectories based on the gradients of the posterior space can produce accurate posterior distributions using far fewer samples than more traditional MCMC methods. The No-U-Turn Sampler (NUTS) adaptively tunes the HMC path for more efficient overall Harmonized adjusted feature values are obtained by the equation:

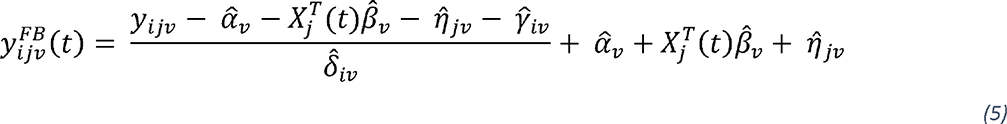

where 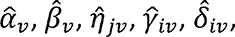 are the posterior parameter estimates. We perform this transformation on the joint posterior parameter distribution to obtain the posterior harmonized data distribution.

### 2.6 Implementation

NUTS inference is implemented in NumPyro Version 0.72 (Phan et al., 2019) using Python version 3.7.1. We run NUTS inference with 4 chains, 40000 samples, and 1000 warmup samples on a CentOS Version 8.2 Linux Machine using 4 Nvidia V-100 32GB GPUs. This inference takes about six hours to complete. Posterior distributions for all parameters are gathered from the 40000 MCMC samples. In both EB and FB ComBat, thickness and volume features are harmonized together.

## 3. Experiments and Results

### 3.1 Overview

We perform several experiments to evaluate the harmonization methods. First, we check HMC sample quality and convergence for the FB ComBat model. Next, we compare harmonization performance in EB ComBat versus FB ComBat with respect to retaining biological (i.e. age and disease) information and removing scanner information. We also explore posterior uncertainty as a tool for dataset augmentation, overall regional measurement uncertainty, and uncertainty-aware association tests between brain regional measures and AD. Next, we perform a simulation study to test how harmonization affects disease effect identifiability. Finally, we perform sensitivity analysis to examine the effect of different priors on FB ComBat harmonization.

### 3.2 Sampling Validation

We check for the quality of our HMC sampling using two methods. We use the Effective Sample Size metric to ensure low auto-correlation of our sampling (Gelman et al., 2021). We also visually inspect the overall likelihood of the model and individual parameter chains to ensure that the inference algorithm converges to the stationary posterior distribution.

Overall model density is shown in Figure 2. After the warmup HMC parameter-tuning phase, all chains converge rapidly and explore the posterior distribution, indicating successful sampling. Effective sample sizes for various parameters are shown in Supplementary Table S1.

**Figure 2:**
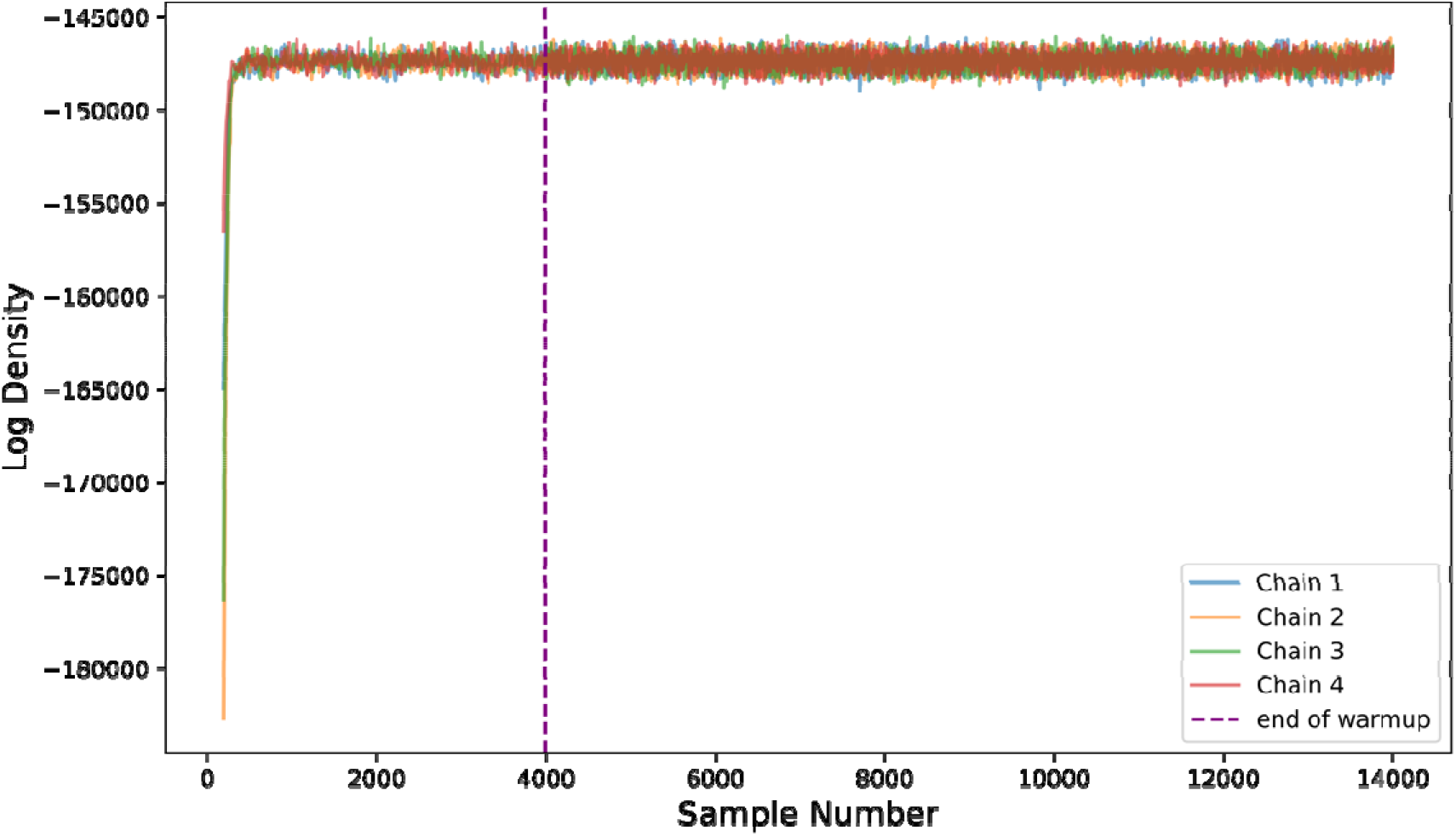
Sampling model density Log of joint density for FB ComBat model given all parameters in each sample. All four chains are shown. The chains converge to the high probability region of the posterior distribution and exhibit good mixing (rapidly exploring the full region), and stationarity.

### 3.3 Posterior Distribution

Posterior distributions for harmonized imaging measurements are obtained for EB ComBat and FB ComBat harmonization. Posterior variances from FB ComBat are larger than those from EB ComBat, as they incorporate uncertainty from all parameters of the ComBat model. An example of a posterior distribution for a single measurement (left entorhinal cortex thickness) from one image is shown in Figure 3. Prior and posterior estimates for individual parameters are shown in Supplementary Figure S1.

**Figure 3:**
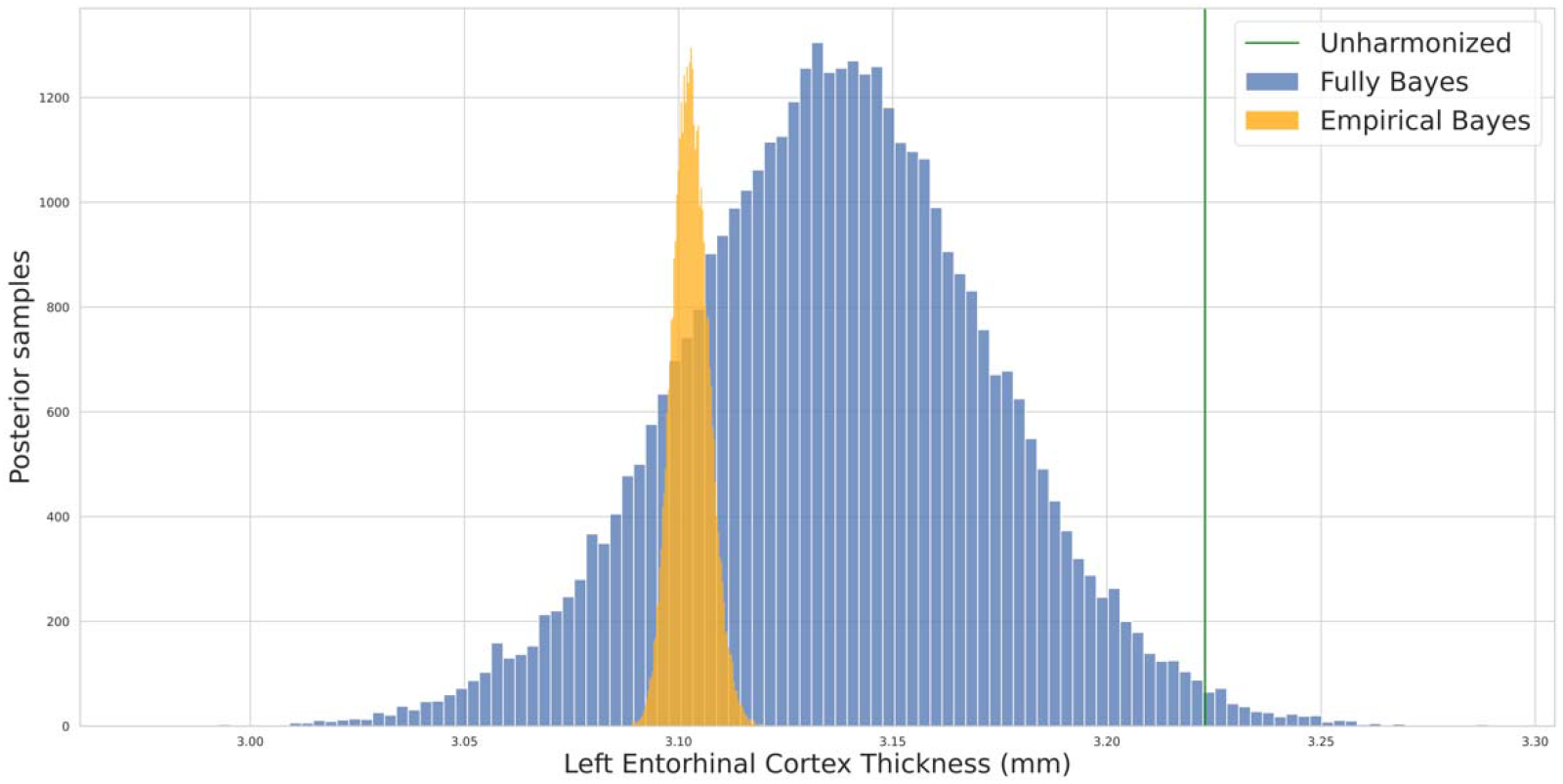
Single Measurement Posteriors Harmonized posterior distribution for left entorhinal cortex thickness from a single image. The FB harmonized posterior is noticeably wider than the EB harmonized posterior. The unharmonized thickness value is shown in green.

### 3.4 Exploratory Analysis of Scanner Effects

We first explore and visualize the unharmonized and harmonized data for the presence of scanner effects. We check for significant additive and multiplicative scanner effects in all features using the Kenwald-Roger (Kenward and Roger, 1997) and Fligner-Killeen (Conover et al., 1981) tests using the longitudinal ComBat package (Beer et al., 2020). Of the 122 imaging features, significant (*p<0.05*) additive effects are present in 121 (of 122) features in unharmonized data and zero features in both EB ComBat-harmonized data and FB ComBat-harmonized data. Multiplicative effects are present in all 122 features in unharmonized data, 1 feature in EB ComBat-harmonized data, and 6 features in FB ComBat-harmonized data.

A visualization of scanner effects (after regressing out covariates and z-score normalization) across field strengths and manufacturers is shown in Figure 4. We plot the distribution of feature values for each image and summarize the distributions across each of the six field strength/manufacturer combinations. The field strength and manufacturer differences from unharmonized data are partially (and to a similar degree) removed to a similar degree by EB and FB ComBat.

**Figure 4:**
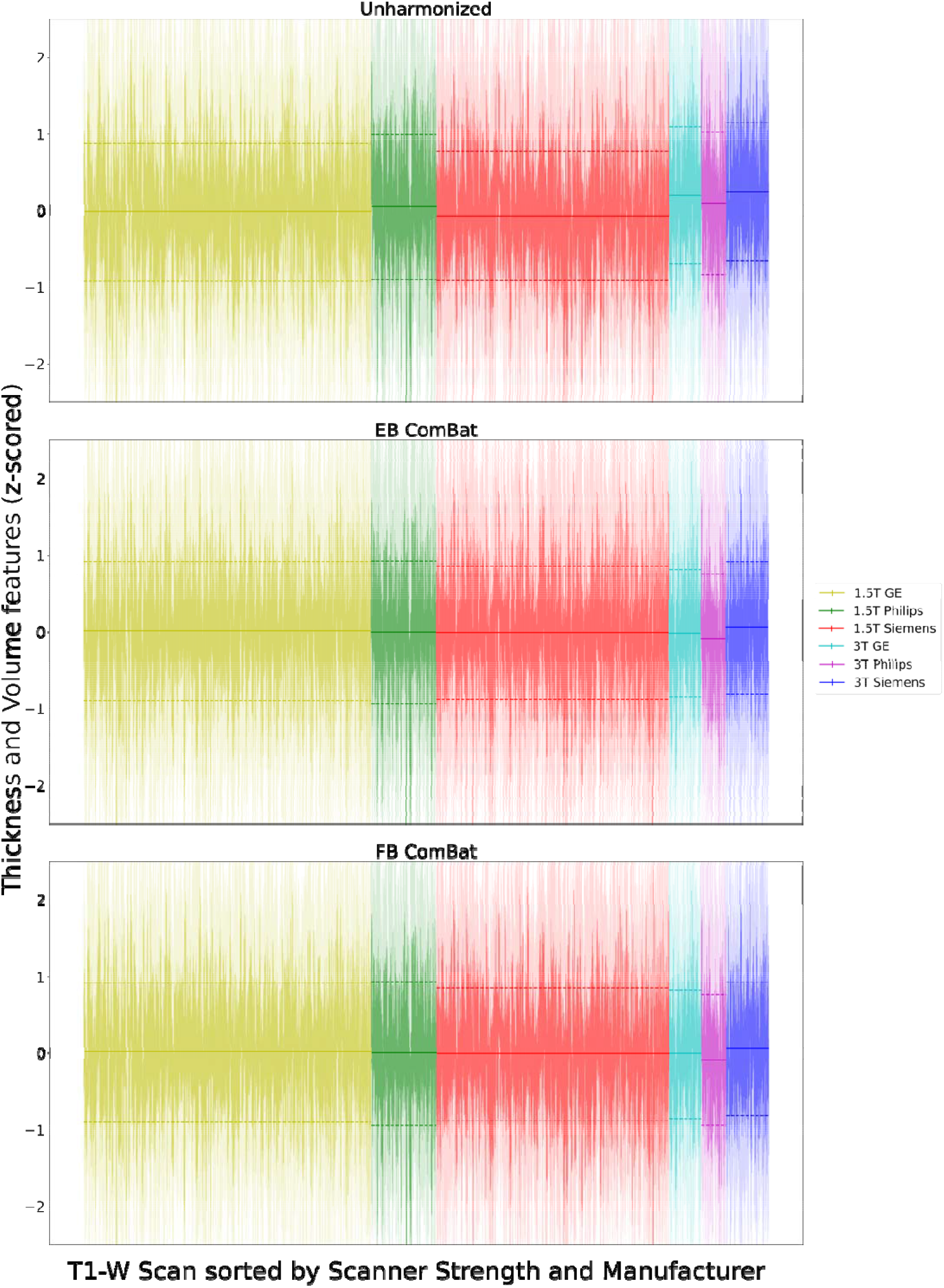
Scanner Effects across Field and Manufacturer Each vertical line represents the distribution of normalized thickness and volume features for a single image; the region within one standard deviation of the image-specific mean is shaded darker. Horizontal lines show the means and 1 standard deviation intervals of normalized feature means for each scanner strength/manufacturer combination. Covariate effects are regressed out before plotting. Distribution distances across field strength and manufacturer are partially but not completely removed by both EB ComBat and FB ComBat.

We also performed linear discriminant analysis (LDA) on the imaging features using the field strength and manufacturer together as the target variable. The first three components in unharmonized, EB ComBat-harmonized, and FB ComBat-harmonized data are shown in Figure 5. Both EB ComBat and FB ComBat remove most variation associated with scanner field strength and manufacturer, although some scanner-related variation is still evident in FB ComBat data (i.e. between 1.5T GE and 1.5T Siemens scanners in the first two LDA components).

**Figure 5:**
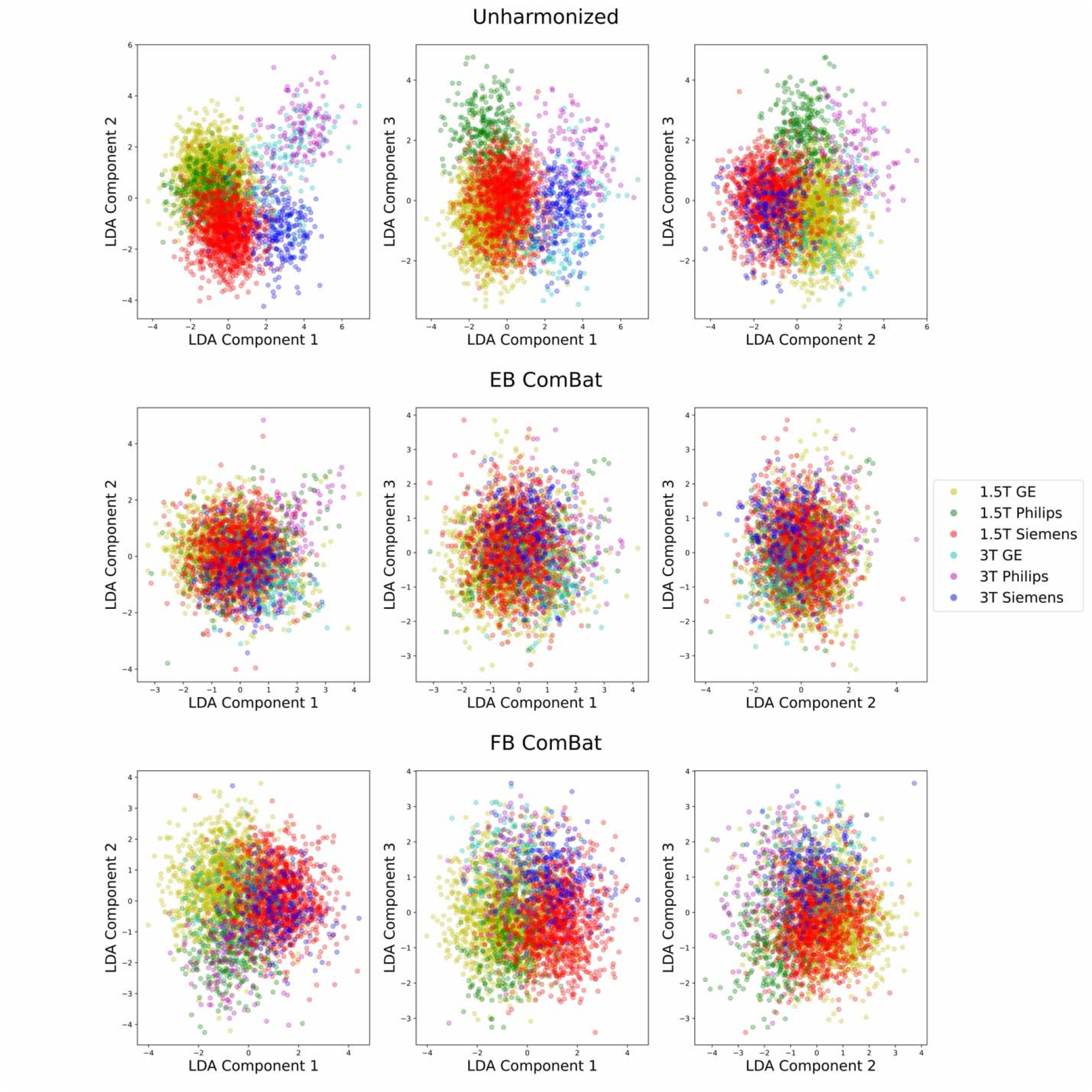
Linear Discriminant Analysis (LDA) with Respect to Field Strength and Manufacturer The first three LDA components of unharmonized and harmonized datasets (using the combination of scanner strength and manufacturer as the target variable) are shown. EB ComBat and FB ComBat remove most scanner-related variation in LDA components, although some scanner-related variation is still visible in FB ComBat data. Covariates are regressed out from features used for LDA, and these features are z-score normalized.

### 3.5 Age Prediction

One objective of harmonization is to retain biological information in the adjustment. To evaluate whether the age information is preserved after harmonization, we train three separate random forest regression models (Breiman, 2001) from all 122 brain measures from 1) unharmonized data, 2) EB-harmonized data, and 3) FB-harmonized data. Age prediction performance of harmonized brain features has been used previously to validate the retention of biological information after harmonization (Fortin et al., 2018; Wachinger et al., 2021). We use three separate sets of predictors: unharmonized measurements, EB ComBat harmonized measurements, and FB ComBat posterior mean harmonized measurements, and compare mean absolute error (MAE) and R^2^ for all three models using repeated k-fold validation with three repeats and 10 folds. We also include a dummy classifier that outputs the mean age value in the training set as a baseline that ignores input. We evaluate performance by using Wilcoxon signed-rank test for cross-validation MAE scores. In the data used for the age prediction task, the mean age is 76.3 years (min = 55 years, max = 93 years) and the data are approximately symmetric (skew = -0.38).

Age prediction results for unharmonized data, EB ComBat, and FB ComBat are shown in Table 2. FB ComBat results in a lower test MAE for age prediction than EB ComBat*(p<10*^-6^), indicating that less age-related biological information is removed in FB ComBat. The test MAE using unharmonized data is not significantly lower than using FB ComBat *(p=0.39)*.

**Table 2:** Age Prediction Results Mean absolute error (MAE) and R^2^ are shown for the age prediction task evaluating retention of biological (age-relevant) information after harmonization. Cross validation standard deviation is shown in parentheses. FB ComBat has a lower test MAE and higher test R^2^ compared with EB ComBat indicating that FB ComBat performs slightly better than EB ComBat.

| Method | MAE (years) | | $R^2$ | |
| --- | --- | --- | --- | --- |
|  | *Train* | *Test* | *Train* | *Test* |
| Dummy Classifier | 5.38 (0.19) | 5.38 (0.19) | 0.00 (0.01) | 0.00 (0.01) |
| Unharmonized | 1.05 (0.01) | 2.79 (0.14) | 0.96 (0.00) | 0.70 (0.02) |
| EB ComBat | 1.09 (0.01) | 2.92 (0.15) | 0.95 (0.00) | 0.67 (0.02) |
| FB ComBat | 1.05 (0.01) | 2.80 (0.13) | 0.96 (0.00) | 0.70 (0.02) |

Differences in population characteristics across scanners may cause identifiability issues for scanner and covariate effects. We therefore check for mean differences in age across scanners using one-way analysis of variance (ANOVA). We check for age differences at two levels: individual scanners (83 groups) and field strength/manufacturer combinations (6 groups). We found significant age mean differences (*p=0.001*) at the individual scanner level but no significant age mean difference (*p=0.501*) at the field strength/manufacturer level.

### 3.6 Scanner Strength Prediction

The harmonization process should remove the effect of non-biological covariates that introduce bias in data. An example of such covariate is variation in the scanner strength. We train random forest binary classifier models using the three datasets (unharmonized, EB harmonized, and FB harmonized) to predict scanner strength (3.0T vs 1.5T). For this task, achieving high accuracy indicates that scanner information remains in the data. In other words, low classifier accuracy is an indicator of better harmonization. We report the area under the receiver operating characteristic curve (AUROC) for evaluation. We use Wilcoxon signed-rank test for cross-validation AUROC scores to assess the statistical significance between the performance of two models.

Area under the receiver operating characteristic curve (AUROC) curves and values of scanner strength prediction from the random forest classification model trained on unharmonized, EB ComBat, and FB ComBat data are shown in Figure 6. Training AUROC values are listed in Supplementary Table S2. Lower accuracy on this task suggests that scanner strength information was more effectively removed using EB ComBat than FB ComBat*(p<10*^-4^), although both methods show improvement over unharmonized data.

**Figure 6:**
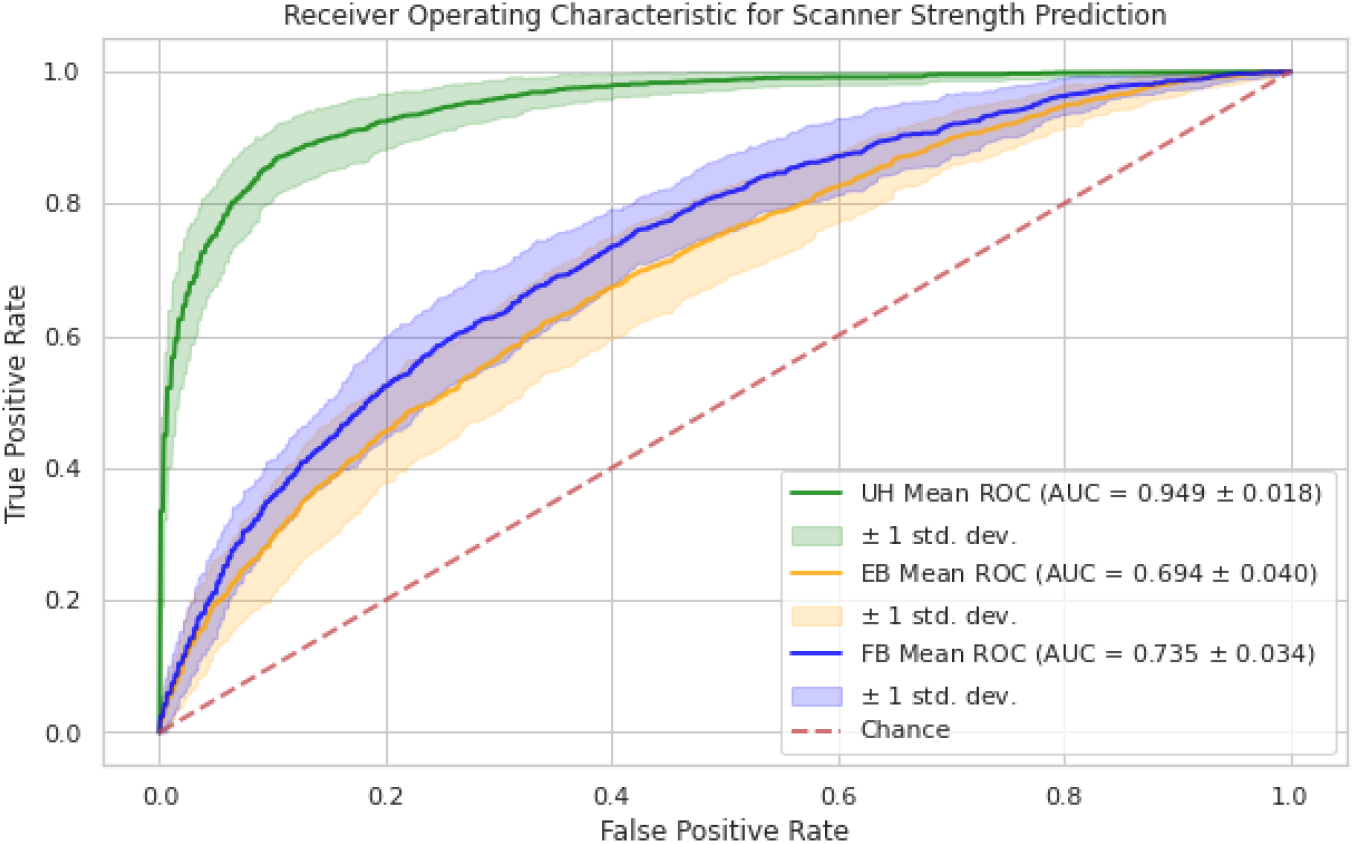
Scanner Strength Prediction AUROC AUROC curves for scanner strength prediction in unharmonized (UH), EB harmonized, and FB harmonized imaging features. Means of cross-validation AUROC are shown by green, orange, and blue lines; coverage envelope of one standard deviation of cross-validation AUROC is shown by colored shaded regions. Lower AUROC indicates better performance for this task. Both EB and FB ComBat similarly reduce the AUROC closer to random chance, but there is still some scanner signal left in the harmonized data.

### 3.7 Test-retest using Paired Scan Evaluation

For the next phase of evaluation, we identified 184 imaging pairs where a subject was scanned on two different scanners on the same day. In all cases, the subject was scanned on both a 1.5T and a 3T scanner. The ground truth difference in brain thickness and volume should be negligible between same-day measurements. Perfect harmonization therefore should remove any difference between these imaging pairs.

Following previous work (Torbati et al., 2021b, 2021a), we compare differences between the paired images on unharmonized, EB harmonized, and FB harmonized datasets for 22 imaging features that have previously been selected as regions of interest for studying AD (Pölsterl and Wachinger, 2020). The mean difference between paired 3T and 1.5T features (bias) and root mean squared deviation (RMSD), a measurement of variance, are computed in the three datasets. We use paired T-tests across the paired images to identify significant bias with respect to scanner strength in any of the three datasets. We also use paired T-tests on the mean absolute differences in image pairs to compare the harmonization performances of EB ComBat and FB ComBat. Significance thresholds for bias and absolute error are adjusted for multiple tests using Bonferroni correction.

Before harmonization, significant biases are found in 16 of 22 regional measurements, shown in Figure 7. Both EB and FB ComBat harmonization remove any significant bias in all regions. RMSD values are shown in Figure 8. All regions had the lowest RMSD after FB ComBat harmonization. Both FB ComBat and EB ComBat improve error variance in all regions compared to unharmonized data. For all thickness and volume measurements, both harmonization methods (EB and FB ComBat) decrease the mean absolute difference between paired scans across scanners, as shown in Figure 9. Additionally, FB ComBat paired scan values have significantly smaller absolute differences (*p<10*^-14^) consistently for all 22 volume and thickness measurements compared to EB Combat.

**Figure 7:**
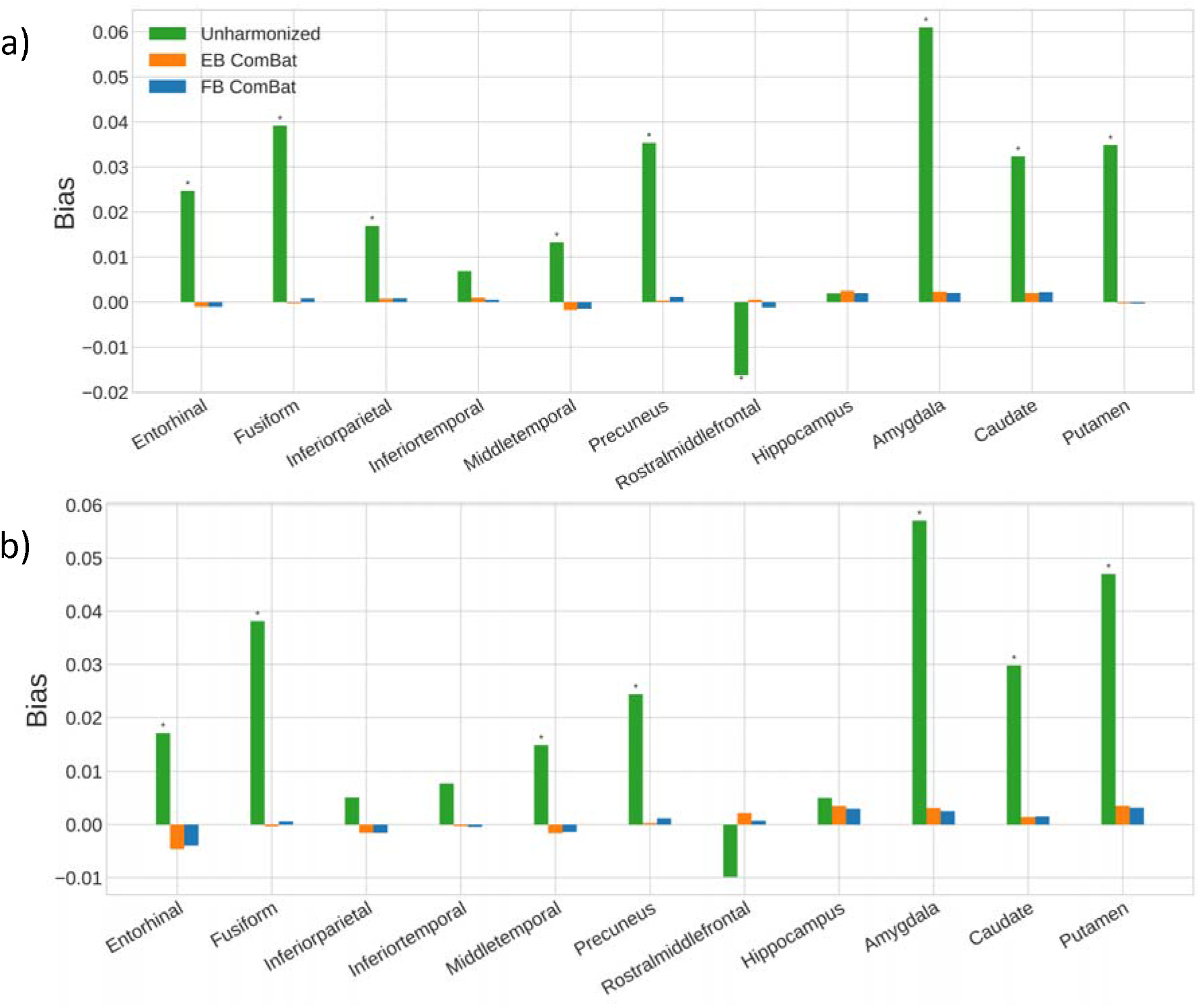
Test-retest Scanner Strength Bias Bias (mean difference) between 3T and 1.5T test-retest scans in AD-relevant brain regions for left hemisphere (a) and right hemisphere (b) measures. Significant biases (p<0.05) are denoted with (*). Significant bias is present in all but six regions for unharmonized data and is removed in all regions with both EB ComBat and FB ComBat. Biases are normalized with respect to the mean feature value.

**Figure 8:**
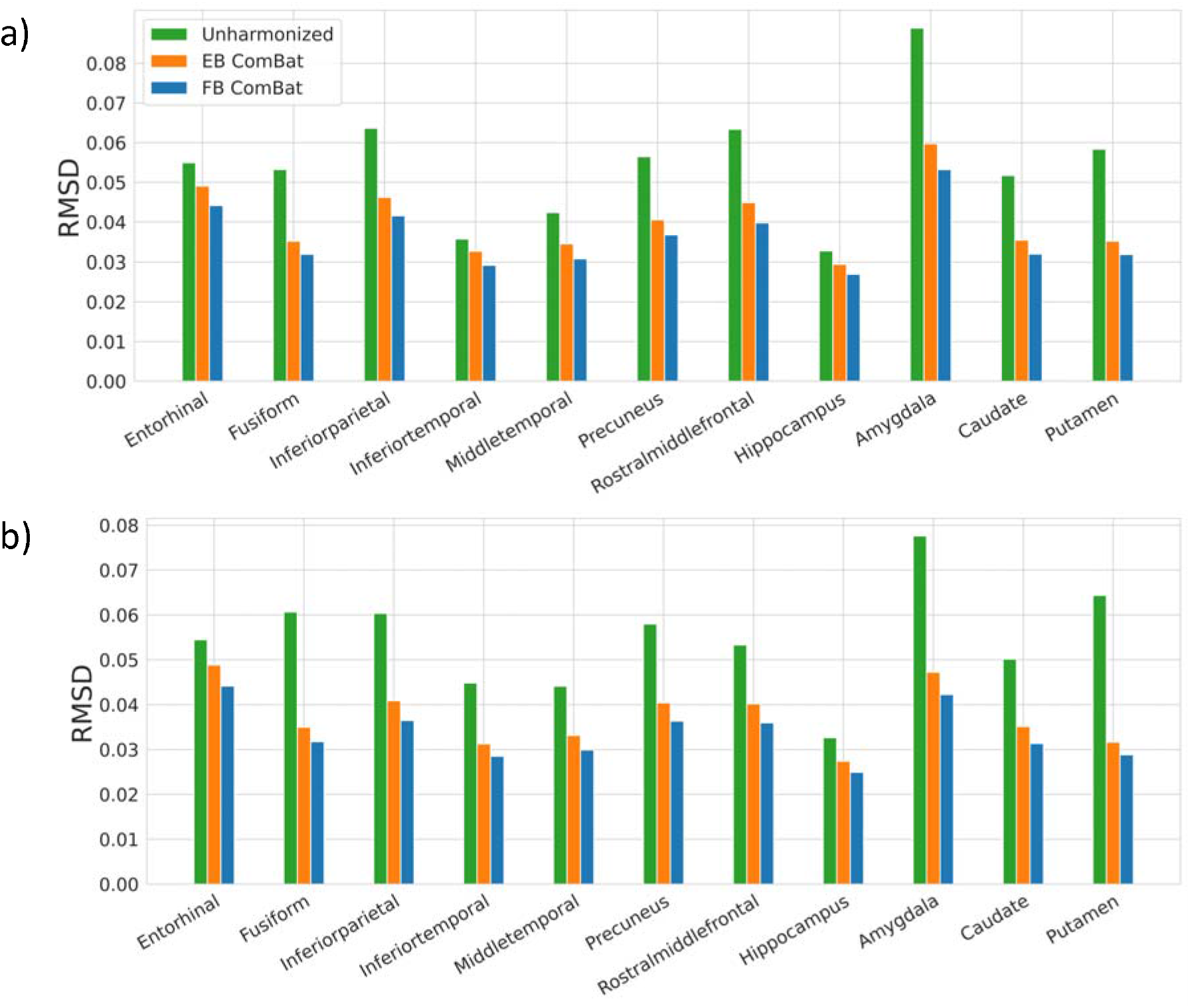
Test-retest Scanner Strength Variance (RMSD) Variance (Root mean squared deviation) between 3T and 1.5T test-retest scans in left hemisphere (a) and right hemisphere (b) AD-relevant brain regions. Variances are normalized with respect to the mean feature value. All regions have the lowest RMSD after FB ComBat harmonization.

**Figure 9:**
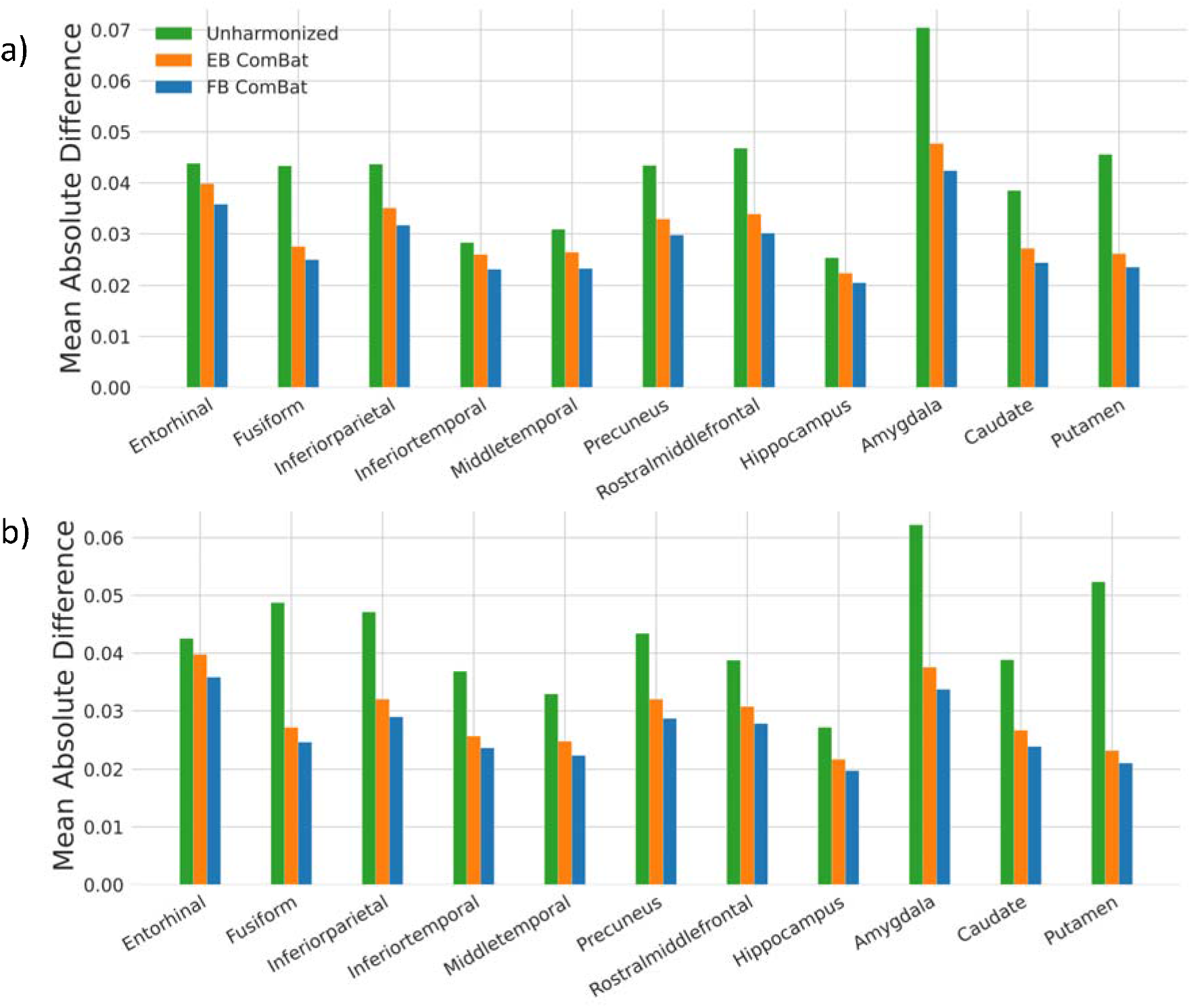
Test-retest Harmonization Error Absolute mean difference in test-retest scans regions for AD-relevant left hemisphere (a) and right hemisphere (b) measures. FB ComBat paired scan values have significantly smaller absolute differences (p<10^-13^) for all 22 volume and thickness measurements compared to EB ComBat, indicating better harmonization performance on the test-retest scans. Errors are normalized with respect to mean feature value.

### 3.8 Dataset Augmentation

Dataset augmentation involves artificially increasing the size of a dataset by modifying the existing data or creating synthetic data. Augmentation is often used to increase classifier performance (Wong et al., 2016). In the imaging domain, augmentation is performed by applying transformations to an image that do not change its content or class label. For tabular data such as FreeSurfer regional thickness and volume features, data augmentation is not as straightforward. We hypothesize that the posterior probability distribution of our harmonized features can be sampled to augment tabular imaging feature datasets. Specifically, each harmonized image can be seen as a *V*-dimensional (corresponding to the number of imaging features) probability distribution. In addition to using the posterior mean of each image, as in typical ComBat harmonization, we also randomly sample from each distribution a number of times. This increases the size of our training set.

We evaluate EB ComBat and FB ComBat as tools for data augmentation by generating additional data samples for a prediction task, namely classifying a patient as having MCI or AD based on imaging features. We perform scanner-stratified train/test splits and repeat 20 times via different random seed numbers. The stratification ensures different scanner groups are used for train and test splits. Images from 75% of scanners are used for training, and images from the remaining 25% of scanners are used for validation. Of the 2408 MCI and AD images from 596 subjects, training sets include between 1609 (66.8%) and 1963 (81.5%) of the images. Five datasets are used in this evaluation. First, the three datasets—unharmonized, EB ComBat, and FB ComBat—are used. Augmented EB ComBat and FB ComBat datasets are then created. For every image in the training split in these two datasets, we draw 100 additional random samples from the posterior distribution of the image’s harmonized thickness and volume features. An augmentation that improves classification performance is desirable. We use the AUROC of AD classification to evaluate the samples generated from EB ComBat and FB ComBat.

Predictive performance for classifying MCI versus AD patients with and without augmentation is shown in Table 3. Without any augmentation, FB ComBat outperformed EB ComBat (*p=0.002*). Using FB ComBat harmonization with posterior distribution resampling augmentation results in the highest AUROC. This is significantly higher than FB harmonization without augmentation (*p<0.0001*) and higher than EB harmonization with augmentation (*p<0.0001*).

**Table 3:** Disease Prediction Results Evaluation of MCI versus AD classification with and without augmentation (sampling from the posterior distribution of harmonized imaging features). Area under the receiver operating characteristic curve (AUROC) is greatest in FB ComBat with augmentation.

|  | **AUROC** |
| --- | --- |
| Unharmonized | 0.772 (0.028) |
| EB ComBat | 0.777 (0.026) |
| FB ComBat | 0.796 (0.030) |
| EB Combat with posterior resampling | 0.799 (0.027) |
| FB ComBat with posterior resampling | 0.821 (0.022) |

As with age (see Section 3.5), understanding scanner distributions of disease diagnosis can reveal a potential source of confounding for scanner and covariate effects. We use chi-squared test of independence to check for an association between scanner and diagnosis. We find no significant association between scanner and diagnosis both at the individual scanner level (*p=0.91*) and field strength/manufacturer level (*p=0.38*).

### 3.9 Region-Level Uncertainty

To identify brain measurements most and least prone to measurement uncertainty, we use the posterior variance of the FB-harmonized imaging features. To compare uncertainty across regions of interest in the brain, it is necessary to normalize posterior variance, due to the difference in measurement scales (thickness vs. volume). Additionally, different regions also have different variances across the population, even after adjusting for mean thickness or volume. We devise a normalized uncertainty value as the ratio of average variance within all *individual* measurements (MSW-Mean Squared Within) to the variance of individual measurement posterior means among the population (MSA-Mean Squared Among) for each regional imaging feature (*v*):

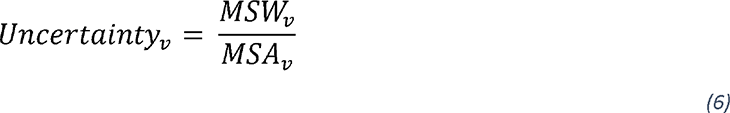

We compute relative uncertainty values for thickness and volume measurements of regions on the Desikan-Killiany cortical atlas (Desikan et al., 2006) and Freesurfer Volumetric Segmentation Atlas (Fischl et al., 2002).

Region-level uncertainty results are shown in Figure 10. Subcortical volumes generally have lower uncertainty values compared to cortical thickness. Among cortical thickness features, regions in the temporal lobe have lower overall uncertainty. Left and right pericalcarine thickness have the highest overall uncertainty among regional imaging features. Among subcortical volume features, mid-anterior and posterior regions of the corpus callosum have the highest posterior uncertainty.

**Figure 10:**
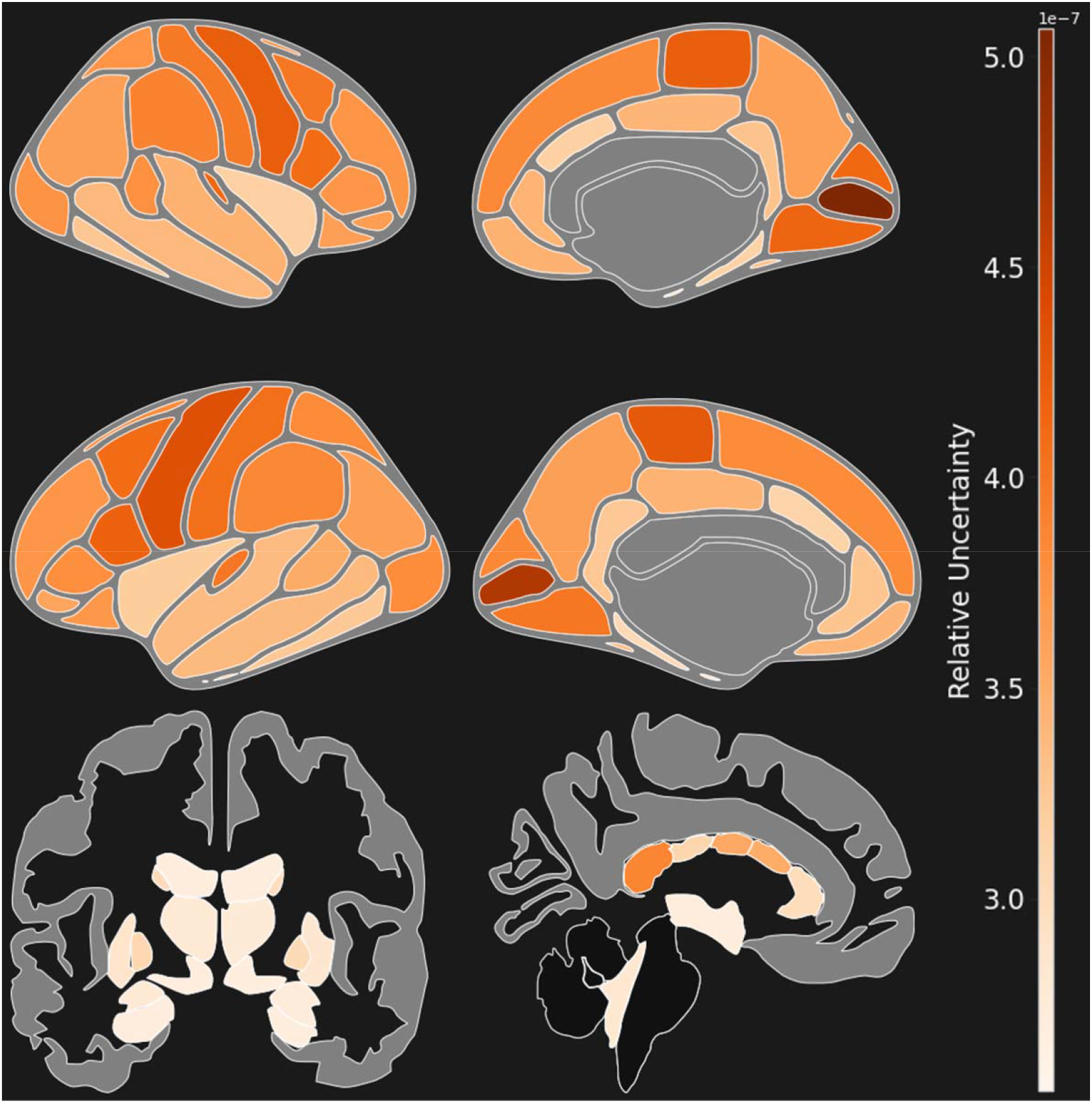
Brain-Wide Uncertainty Regional uncertainty, defined as the ratio of variance within the posterior distribution of individual measurements (MSW-Mean Squared Within) to the variance among the population (MSA-Mean Squared Among). Uncertainty is shown in cortical thicknesses and subcortical volumes. Uncertainty is higher in cortical thickness values compared to subcortical volumes. Left and right pericalcarine thickness have the highest overall uncertainty of any region.

### 3.10 Uncertainty in Statistical Analysis

Statistical association models like least squares regression, which can be used to model brain regional associations with disease (Wang et al., 2011), consider all data points as equally reliable when minimizing the error term. In the brain imaging domain, this assumption may not hold when scanner reliability introduces uncertainty in a multi-site study. We propose using the posterior variance of harmonized measurements as a measure of uncertainty. Data for a regression model can then be weighted by the inverse of this variance, resulting in a model fit that accounts for measurement uncertainty (Aitken, 1936).

We demonstrate the difference between ordinary least squares (OLS) and uncertainty-weighted least squares (WLS) by testing for association between different brain region measurements with Alzheimer’s disease. We use the model:

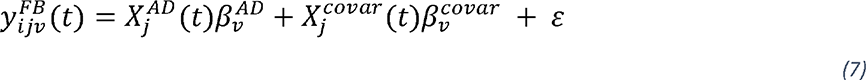

with null and alternative hypotheses:

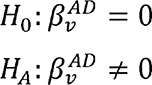

where 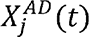 is an indicator variable for whether a patient has an Alzheimer’s disease diagnosis at time *t*, 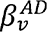 is the association of Alzheimer’s disease with feature *v*, 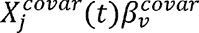 is the covariate term, and e is the random error term. Imaging features 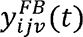 are adjusted by mean thickness or intracranial volume for regional thickness and regional volume measurements, respectively. A significant 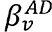 indicates an association between the feature *v* and Alzheimer’s disease, adjusting for covariates (age and sex).

We test for association between 104 brain regions (from Desikan-Killiany and Freesurfer Volumetric Segmentation atlases) and Alzheimer’s disease using OLS (baseline) and WLS weighted by the reciprocal of measurement posterior variance and examine differences in findings between the two methods.

Significant associations (after Bonferroni correction) between brain regions and Alzheimer’s disease are shown in Figure 11. Several regions show differing significance when using WLS versus OLS regression including right lateral occipital thickness, left rostral anterior cingulate thickness, and left caudate volume. We also show results from using EB ComBat posterior variance in Supplementary Figures S2,S3.

**Figure 11:**
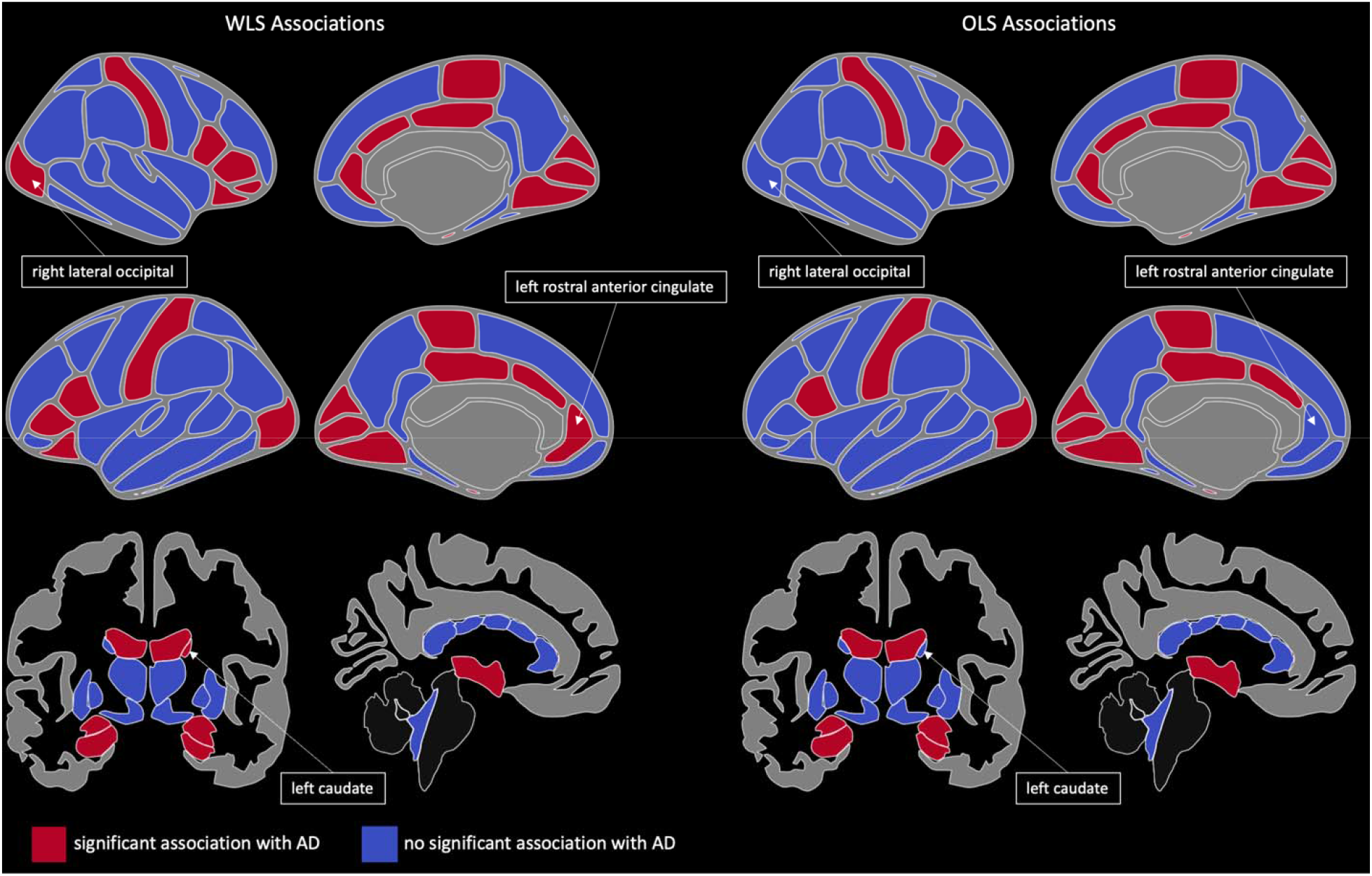
Cortical Thickness and Subcortical Volume Associations with AD Cortical thickness and subcortical volume regions associated with Alzheimer’s disease using Ordinary Least Squares (OLS) and Weighted Least Squares (WLS) weighted by the reciprocal of posterior measurement variance. Right lateral occipital, left rostral anterior cingulate thickness, and left caudate volume are highlighted as regions with different associations using OLS and WLS models.

### 3.11 Simulation Study

Following prior work (Beer et al., 2020), we conducted a simulation analysis to study to impact of harmonization on the ability to identify disease effects on neuroimaging features. First, we segregate the AD and CN patients from ADNI and fit the following linear mixed-effect model to each imaging feature separately:

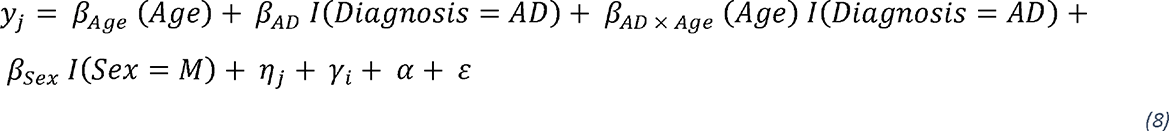

where *y_j_* is the feature value for subject *j*, *β* are coefficients for covariates (age, AD, age × AD interaction, and sex), *η_j_* is a random subject-specific intercept, *γ_i_* is a scanner-specific intercept, *α* is a feature intercept, and *ε* is the residual. We select six features (right hippocampus volume, left parahippocampal thickness, right accumbens area volume, left medial orbitofrontal thickness, right amygdala volume, and left thalamus volume) with significant *β_AD_* and *β_AD × Age_* coefficient values and choose these as “alternative hypothesis features”. The remaining features are considered null features.

We then isolate the CN ADNI patients, randomly assign half to the AD group, manually add the identified AD and AD × age effects to the AD group patients, and harmonize the semi-synthetic dataset using EB ComBat and FB ComBat. We repeat the process of random group assignment and harmonization 500 times. Finally, we re-fit the linear mixed effect model (without the scanner additive term) on all 500 sets of unharmonized, EB ComBat-harmonized, and FB ComBat-harmonized data.

We are particularly interested in the ability to recover the AD effect and AD × Age interaction effect from alternative hypothesis features while minimizing Type I Error from the null features. Thus *β_AD_* and *β_AD × Age_* estimates from harmonized data should be near their true values; coefficient p-values for null features should be insignificant, and coefficient p-values for alternative hypothesis features should be significant.

FB ComBat generally reduces type I error and improves coefficient estimates for null features compared to EB ComBat; both methods still underperform in most null feature evaluations compared to unharmonized data. Estimated coefficient values for *β_AD_* and *β_AD × Age_* in null features are shown in Figure 12a. Unharmonized data yields the lowest *β_AD_* mean absolute error for null features, while EB ComBat results in greater *β_AD_* error compared to FB ComBat (*p=0.008*). No difference between *β_AD × Age_* error in EB ComBat and FB ComBat for null features is found (*p=0.28*), while EB and FB ComBat both have lower *β_AD × Age_* error compared to unharmonized data (*p<0.004*). *β_AD_* and *β_AD × Age_* Type I error rates for null features are shown in Figure 12b. Unharmonized null features have a significantly lower Type I error rate for *β_AD_* and *β_AD × Age_* compared to EB and FB ComBat (*p<10*^-5^). *β_AD_* Type I error rate is greater in EB ComBat than FB ComBat (*p<10*^-20^); no significant difference for *β_AD × Age_* error rate is found between EB ComBat and FB ComBat (*p=0.24)*.

**Figure 12:**
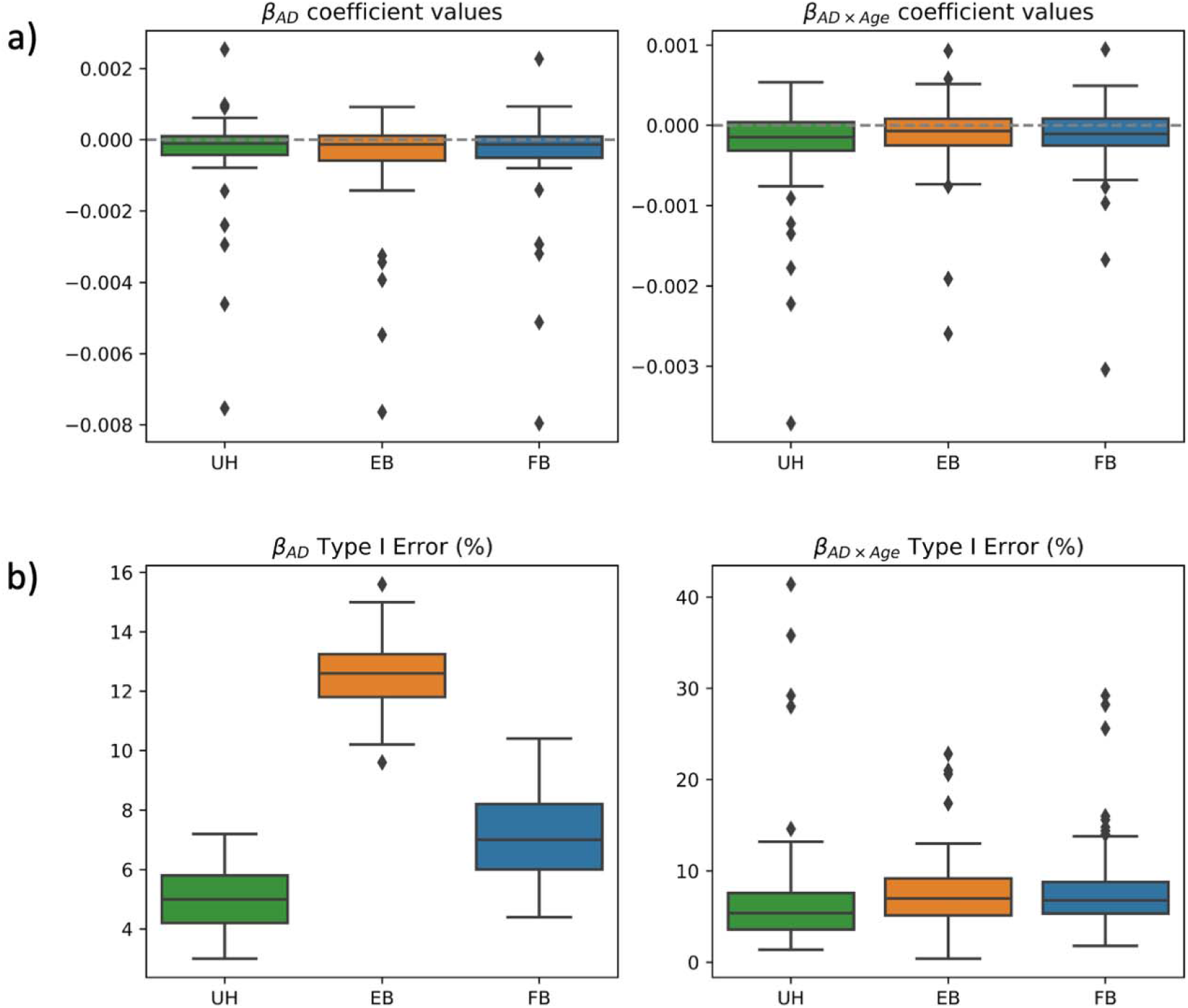
Simulation Study Null Feature Coefficients Simulation data a) Coefficient values and b) Type I error rate for null features (zero AD and AD × Age effect sizes) from unharmonized, EB ComBat, and FB ComBat data. Coefficient values in a) are shown across all null features and simulations and are normalized with respect to feature means. Type I Error rate is calculated as the percentage of Type I error in 500 simulations for a given feature.

EB ComBat leads to the highest sensitivity to *β_AD_* for alternative hypothesis features, while FB ComBat data has the highest sensitivity to *β_AD × Age_* interaction effects. Error in coefficient value estimates for the six alternative hypothesis features do not significantly differ across harmonization measures. Coefficient estimates are shown in Supplementary Figure S1. P-value distributions for these features are shown in Figure 13. The *β_AD_* p-values for all six features are more significant in EB ComBat than FB ComBat (*p<10*^-4^) and are more significant in both EB and FB ComBat compared to unharmonized data (*p<10*^-5^). The *β_AD × Age_* p-values for five of six features are more significant in FB ComBat than EB ComBat (*p<0.01*). All *β_AD × Age_* p-values are more significant in FB ComBat than in unharmonized data, and four of six p-values are more significant in EB ComBat than in unharmonized data.

**Figure 13:**
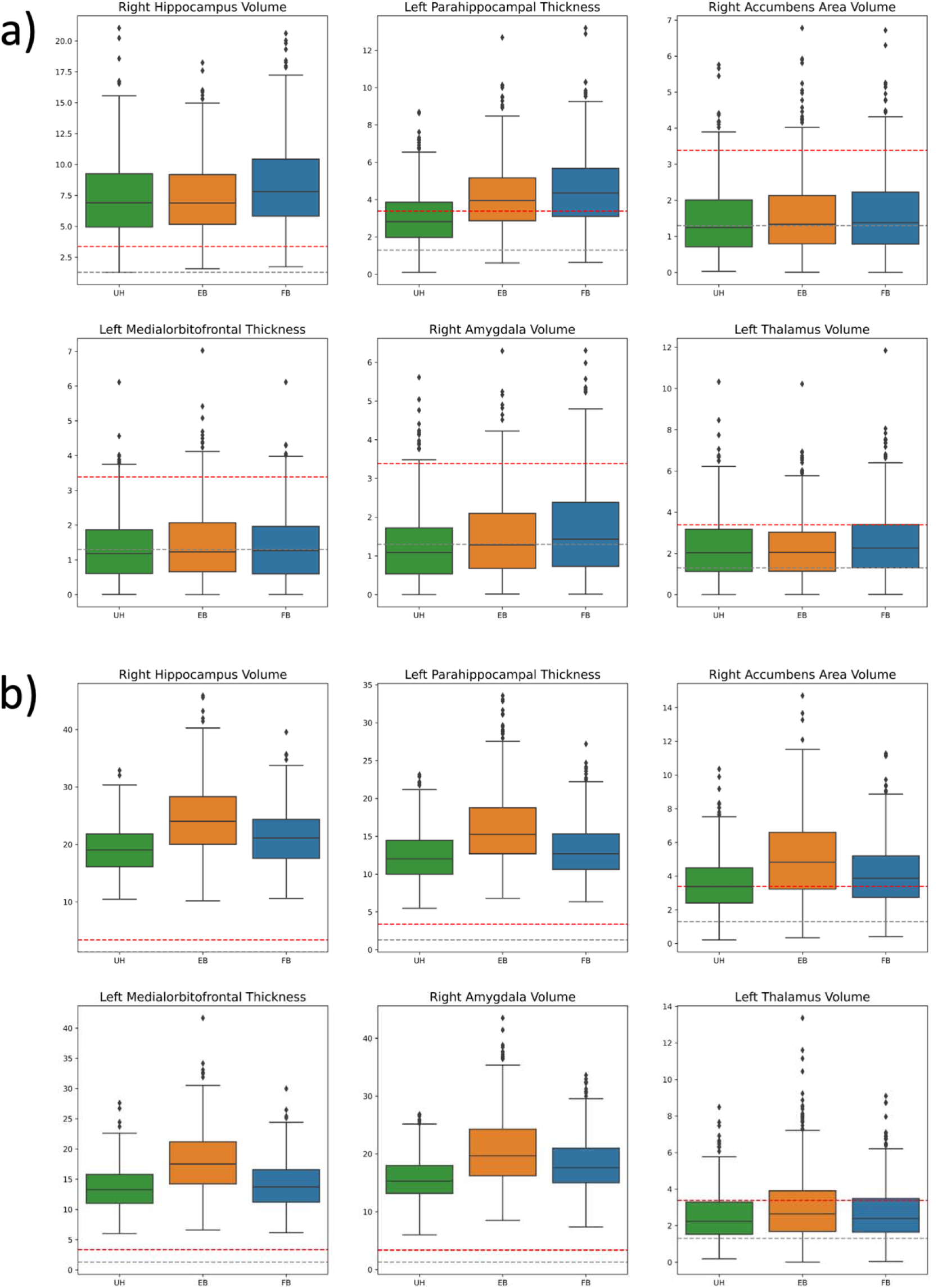
Simulation p-value Disease Effect Distribution Distribution of -log_10_ p-values in the true (alternative hypothesis) disease effect features for a) AD coefficient *β_AD_* and b) AD × Age coefficient *β_AD × Age_*. The dotted grey line is at p=0.05, and the dotted red line is at the Bonferroni-corrected significant p-value (p=0.0004).

### 3.12 Sensitivity Analysis

We perform sensitivity analysis to explore the impact of prior distributions on harmonization. In one set of experiments, we vary the prior distributions *τ_i_* parameter to study whether a strong prior on the presence or absence of scanner effect pooling affects the harmonization performance. We also check whether making the baseline FB ComBat priors stronger changes harmonization. A more in-depth description of our sensitivity analysis procedure can be found in Supplementary Section S4. Overall, we find that harmonization performance was fairly robust to various priors over the hyperparameters. Of note, a strong prior against pooling (high *τ_i_*) leads to modest increases in significant additive scanner effects after harmonization. Stronger priors on all parameters, compared with weakly informative priors, lead to increases in significant multiplicative scanner effects after harmonization.

## 4. Discussion

Scanner harmonization, especially for image-derived thickness and volume features, is an important step of brain MRI pre-processing to reduce noise and potential biases. We built on ComBat, a popular statistical harmonization approach, by proposing a new Fully Bayes method for inference which leads to improved harmonization and additional ComBat use cases. While the FB method maintained the robustness of the EB approach, we also found that FB ComBat yielded harmonized features with greater retention of biologically relevant information and smaller differences in test-retest subjects. EB ComBat, on the other hand, removed scanner strength information more effectively and takes only a fraction of the runtime compared to FB ComBat. FB ComBat produced more realistic posterior distributions and uncertainty quantification, which is important for individualized disease diagnosis (Liu et al., 2020) and group-level statistical analysis (Aitken, 1936). Fully Bayesian inference allows us to draw samples from a rich posterior distribution, which we used to augment a dataset to improve Alzheimer’s Disease classifier performance. We also used the variance of the posterior distributions to perform a more principled analysis of brain regional associations with Alzheimer’s disease (AD) and identify features that are more prone to measurement uncertainty.

Two important metrics of harmonization are that, first, biological information is maintained after harmonization and, second, that scanner-related information and other confounding nuisance are successfully removed. To evaluate EB and FB ComBat’s ability to retain biological information, we trained models to predict age and AD status, and we evaluated their performance on datasets with unharmonized versus harmonized data. Harmonizing brain imaging feature data using FB ComBat resulted in stronger age and AD classification performance than EB ComBat, suggesting that FB ComBat was able to more effectively retain biologically-relevant brain structural information. Our age regression findings align with a recent study finding that ComBat harmonization methods tend to either modestly obscure or not affect biological effects compared to unharmonized data (Richter et al., 2022). Conversely, our AD classification study showed that FB ComBat improved disease identifiability. Richter et al. (2022) designed a more prospective study where subjects were scanned at an initial time point, then several years later at both the initial scanner and a separate scanner, allowing direct comparison of imaging feature change over time with separate scanners compared to a ground truth change. FB ComBat should be evaluated in a similar design against other ComBat approaches to further study its impact on biological and non-biological effect identifiability.

We also trained a model to predict scanner strength from harmonized imaging features. In this task, lower classification accuracy was ideal, indicating that information about scanner strength was removed during harmonization. EB ComBat performed best on this task, indicating that it removed more non-biological information related to scanner strength compared to FB ComBat. Both EB ComBat and FB ComBat also similarly removed detectable effects from scanner strength and manufacturer. Furthermore, we showed that the harmonization performance of FB ComBat was very stable when varying the prior distributions.

One concern with covariate and scanner prediction on imaging features is the potential confounding of covariate and scanner effects. This complicates the identifiability of covariate and scanner model parameters, which can lead to the inadvertent removal of covariate effects or preservation of scanner effects. Additionally, when predicting covariates to evaluate harmonization, models could be learning scanner effects. Models predicting scanner strength may also be capturing covariate effects. Our ANOVA and chi-squared tests found that, among scanner strength and manufacturer, two possible contributors to scanner effect differences (Han et al., 2006), no significant disease or age differences exist. However, some age difference was found across individual scanners. This indicates the possibility that scanner effects related to some factor besides magnetic field strength or manufacturer could be learned by an age prediction model. Due to a large number of scanners, pairwise follow-up group comparisons to examine the extent of age differences were infeasible. Future large-scale imaging studies should ensure that covariates (including diagnostic groups) are balanced across scanners so that scanner and covariate effect confounding is eliminated. Furthermore, in studies with fewer sites, traveling subject-based study design and inferring the ComBat model on only traveling subjects can remove the impact of site-based sampling bias and improve harmonization performance (Maikusa et al., 2021; Yamashita et al., 2019). Additionally, researchers using multiple datasets should carefully consider population differences when pooling and harmonizing the datasets together.

We checked whether the harmonization methods would bring same-day, same-subject imaging features from different scanners closer together. This metric should be seen as a “ground-truth” check, as brain anatomy should not change in such a short time. Our FB ComBat harmonization resulted in the largest reduction in imaging feature measurement difference between repeat scans for all tested regions, indicating that our model specification removed scanner artifact most effectively for this repeat-scan subset of patients.

We further performed a simulation study to determine how both harmonization methods affect the identifiability of disease signals in brain imaging features. Both methods modestly improved identifiability of true effects. EB ComBat led to highest sensitivity in AD effect while FB ComBat led to highest sensitivity in AD × Age interaction effect. We also found that, while both methods increased type I error rate compared to unharmonized data, EB ComBat led to a type I error rate inflation in AD effect estimation compared to FB ComBat. The AD × Age interaction term did not have as pronounced type I error rate inflation, similar to prior simulation studies of EB ComBat (Beer et al., 2020). Further research is still needed to determine strategies for mitigating type I error in harmonization. One potential approach may be to only apply the harmonization transformation when a chosen credible interval (e.g. 95%) of a scanner error term lies above or below 0 (for additive error) or 1 (for multiplicative error). We leave this approach and additional exploration of type I error mitigation for future research.

We also presented the novel use of generative harmonization models to make a downstream classification model more robust with respect to limited training data. Large-scale imaging datasets have grown but are still relatively small in the medical field due to acquisition costs and privacy concerns. Datasets may not contain sufficient variation to train robust classifiers without augmentation. Several studies have examined the effect of image-level augmentations such as random affine transformations, elastic deformation, intensity shifts, or GAN-based image generation on voxel-level tasks such as tumor segmentation or disease classification (Dufumier et al., 2021; Li et al., 2020; Nalepa et al., 2019). However, as far as we are aware, augmentation of regional thickness and volume measurements has not been studied. We presented a new augmentation method that uses the EB and FB harmonization models’ posterior distributions for a rich data generation tool that can improve classifier performance. Our augmentation method draws from our post-harmonization uncertainty regarding measurement error of an image. While EB harmonization generates a posterior distribution, it underestimates posterior uncertainty. Sampling from the EB ComBat posterior for augmentation produces data with less variation, which may explain why FB ComBat performed better than EB ComBat in the augmentation task.

We explored posterior distributions to determine which imaging regions have the most uncertainty, compared to overall population variance. Subcortical volume measurements and temporal thickness were generally less prone to uncertainty than other cortical thickness measurements. The difference in uncertainty between cortical thickness versus structural volumes may be due to the difficulty of surface parcellation compared to segmentation. Gyral-based parcellation, used in FreeSurfer, is inherently difficult because gyri are connected without a clear visible boundary between connected regions (Meng et al., 2015). Our results suggest that subcortical volume measurements may be more reliable than cortical thickness, a finding verified by recent test-retest analysis of Freesurfer measures (Hedges et al., 2022).

Finally, we propose the use of uncertainty-based measurement weighting in association tests. Commonly used models such as ordinary least squares assume that regressors have no error and response variables have uniform uncertainty. We demonstrated that tests involving regional association with Alzheimer’s Disease can vary depending on whether uncertainty is considered (using weighted least squares regression), or it is ignored (using ordinary least squares regression). FB ComBat’s uncertainty measurement should be used for principled downstream statistical analysis. EB ComBat underestimates uncertainty, so we do not suggest using it for uncertainty-aware downstream tasks. Further work might investigate uncertainty-aware models for predictive tasks such as Alzheimer’s disease conversion. Causal discovery that incorporates measurement error (Zhang et al., 2017) is another potential area for further exploration that may benefit from using FB ComBat measurement uncertainty. Further work might also investigate the impact of different experimental settings, such as the number of features included in FB ComBat harmonization, on posterior measurement uncertainty.

We suggest two important practical guidelines for the use of FB ComBat. First, imaging features should be standardized (i.e. z-scored) before inference time. This prevents misspecification in the hierarchical model parameters which assume that scanner effects derive from a common distribution. Conversely, EB ComBat performs standardization based on fixed effect estimates from the first stage of inference so standardization before harmonization is not necessary. Next, we encourage the use of weakly informative priors, especially in scanner-specific parameters *μ_i_, τ_i_, λ_i_, θ_i_*. For example, the parameter *τ_i_* might use a distribution like *InverseGamma(2,0.5)*. This constrains 91% of the prior density to within 0 and 1. In other words, we allow for a 9% prior probability that the standard deviation of additive effects within a scanner varies by more than one feature-standard deviation. The use of weakly informative prior distributions makes the MCMC sampler more efficient while having a minimal impact on the posterior distribution.

Several limitations exist for ComBat harmonization, and for large-scale Bayesian inference. Harmonization evaluation is inherently limited because ground truth imaging feature measurements are unknown. Developing a quantitative metric to evaluate harmonization algorithms is challenging. Test-retest experiments where traveling subjects are scanned on two scanners in a short time, as well as simulation studies where known effects are added to patient subsets, are our best ground-truth tools for evaluating harmonization. Metrics for explicitly studying the retention of biological information and removal of scanner information are also informative and include tasks like age prediction and scanner strength prediction. However, confounding between biological and scanner factors may limit the usefulness of these metrics. Additionally, these tasks are only approximations and do not include all possible biological and non-biological variables of interest, many of which are unobserved. Including additional covariates in the ComBat model, such as total intracranial volume, may improve harmonization performance (Pomponio et al., 2020). Additionally, the linear nature of our model may leave out important non-linear covariate and scanner effects. Extensions of EB ComBat have shown improved performance by modeling scanner covariance effects (Chen et al., 2021) and non-linear covariate effects (Pomponio et al., 2020). While our work compares EB and FB approaches to Longitudinal ComBat (Beer et al., 2020), more complicated FB models should be compared against their corresponding EB ComBat extensions. These extensions are straightforward to implement in the FB setting; the probabilistic programming approach used for FB ComBat just requires specification of the model, then sampling is done automatically. FB ComBat should also be studied for other imaging features that have benefitted from ComBat harmonization such as PET standardized uptake values (Orlhac et al., 2018), functional MRI (Yu et al., 2018), and white matter imaging features such as fractional anisotropy, mean diffusivity and regional volumes (Fortin et al., 2017; Richter et al., 2022). Finally, FB inference is inherently slower than an EB approach. EB ComBat uses expectation-maximization optimization while FB ComBat relies on much slower MCMC sampling. With large models like ComBat, the difference in inference speed is significant. If speed or computational resources are a concern (e.g. if datasets are constantly updated and re-harmonized), EB ComBat may be preferred. However, harmonization is generally performed just once before imaging features are analyzed, so we expect that in many cases the uncertainty quantification and improved harmonization performance for most metrics outweigh the efficiency drawback.

## 5. Conclusion

We have compared EB and FB approaches to ComBat brain MRI feature harmonization. FB harmonization performed slightly better in most harmonization tasks. We also demonstrated that the posterior distributions of FB harmonized data should be used for any study where the accurate estimation of uncertainty is important. We provided three examples, namely data augmentation, association tests, and brain-wide feature uncertainty quantification, which utilize the posterior distribution given by FB ComBat. The code for FB ComBat is available at *https://github.com/batmanlab/BayesComBat*.

## Acknowledgements

This work was supported by the Pennsylvania Department of Health [grant number 4100087331], National Institutes of Health [grant numbers R01HL141813, 1R01-AG063752, 5T15LM007059], the National Science Foundation [grant number 1839332], Tripod+X, and the SAP SE. This work used the Bridges-2 system, which is supported by NSF award number OAC-1928147 at the Pittsburgh Supercomputing Center (PSC). Data collection and sharing for this project was funded by the Alzheimer’s Disease Neuroimaging Initiative (ADNI) (National Institutes of Health Grant U01 AG024904) and DOD ADNI (Department of Defense award number W81XWH-12-2-0012). ADNI is funded by the National Institute on Aging, the National Institute of Biomedical Imaging and Bioengineering, and through generous contributions from the following: AbbVie, Alzheimer’s Association; Alzheimer’s Drug Discovery Foundation; Araclon Biotech; BioClinica, Inc.; Biogen; Bristol-Myers Squibb Company; CereSpir, Inc.; Cogstate; Eisai Inc.; Elan Pharmaceuticals, Inc.; Eli Lilly and Company; EuroImmun; F. Hoffmann-La Roche Ltd and its affiliated company Genentech, Inc.; Fujirebio; GE Healthcare; IXICO Ltd.; Janssen Alzheimer Immunotherapy Research & Development, LLC.; Johnson & Johnson Pharmaceutical Research & Development LLC.; Lumosity; Lundbeck; Merck & Co., Inc.; Meso Scale Diagnostics, LLC.; NeuroRx Research; Neurotrack Technologies; Novartis Pharmaceuticals Corporation; Pfizer Inc.; Piramal Imaging; Servier; Takeda Pharmaceutical Company; and Transition Therapeutics. The Canadian Institutes of Health Research is providing funds to support ADNI clinical sites in Canada. Private sector contributions are facilitated by the Foundation for the National Institutes of Health (www.fnih.org). The grantee organization is the Northern California Institute for Research and Education, and the study is coordinated by the Alzheimer’s Therapeutic Research Institute at the University of Southern California. ADNI data are disseminated by the Laboratory for Neuro Imaging at the University of Southern California.

