## Supplemental Materials for "ComBat Harmonization: Empirical Bayes versus Fully Bayes Approaches"

***Section S1: Choice of Prior Distributions***

The entire FB ComBat model specification is shown in Figure 1. We discuss the rationale for prior distributions over the highest-level parameters here. In general, we select “weakly informative” priors in order to inject minimal bias into our models while making sampling from the posterior distribution computationally feasible.

$$\mu_{i} \sim C\left( 0,0.1 \right)$$

We choose a Cauchy distribution centered at 0 with scale 0.1 for the prior over the additive scanner error hyperparameter. This fat-tail distribution assumes that most scanners exhibit a relatively small additive effect while leaving the possibility for outlier scanners with a greater effect.

$$\tau_{i} \sim IG(2,0.5)$$

Wide prior distribution (contstrained to positive numbers) over additive effect pooling parameter.

$\lambda_{i} \sim G(50,50)$, $\theta_{i} \sim G\left( 50,1 \right)$, $\delta_{iv} \sim G(\lambda_{i}\theta_{i},\theta_{i})$

The hyperparameters for the variance scaling term $\delta_{iv}$are again weakly informative and constrained to positive numbers. The formulation for $\delta_{iv}$ $G(\lambda_{i}\theta_{i},\theta_{i})$is intended to be interpretable in nature. The $\delta_{iv}$ prior mean is $\lambda_{i}$, and the $\delta_{iv}$ prior variance is controlled by $\theta_{i}$.

$$\rho_{v} \sim IG(6, 5)$$

A weakly informative prior with expectation of 1 on for the variance hyperparameter for subject-specific intercept $\eta_{jv}$.

$$\sigma_{v} \sim HC(0.2)$$

Weakly informative fat-tailed variance hyperparameter, constrained to positive values.

$$a_{v} \sim C(0,0.5)$$

Weakly informative fat-tailed mean hyperparameter centered at zero. Less regularization is needed compared to $\mu_{i} .$

$${p(\beta}_{v})\propto1$$

A uniform distribution is used for covariate terms. We found this did not inhibit sampling efficiency.

***Section S2: Sampling Quality***

| Parameter Type | # of Paramaters | Minimum ESS | Mean ESS | Maximum ESS |
| --- | --- | --- | --- | --- |
| $\alpha$ | 122 | 1511 | 2047 | 2857 |
| $\beta$ | 732 | 295 | 2287 | 4302 |
| $\gamma$ | 10,126 | 719 | 3711 | 17335 |
| $\delta$ | 10,126 | 10772 | 22290 | 28252 |
| $\eta$ | 98,698 | 496 | 5786 | 25955 |
| $\sigma$ | 122 | 13067 | 29796 | 39075 |

*Table S1: Effective sample size (ESS) for parameters in FB ComBat model. Note that there are varying numbers of each parameter type (e.g. for* $\gamma$ there is one additive factor for every scanner *i* for every feature type *v*).

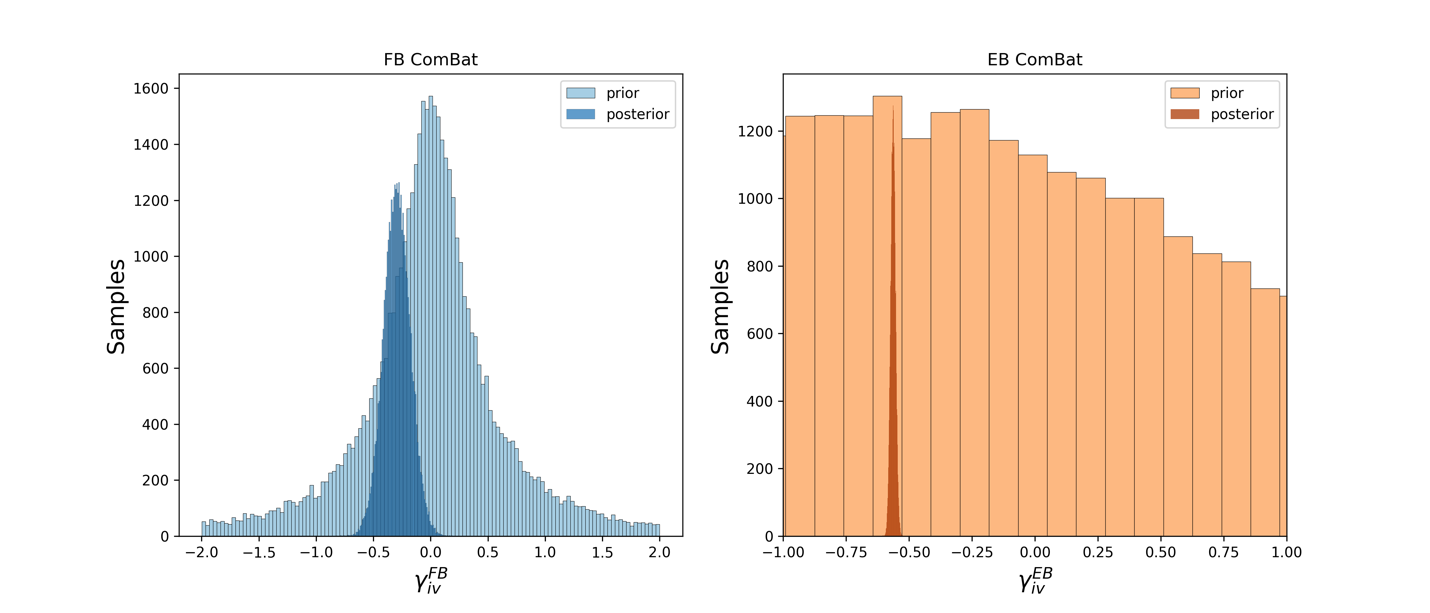

*Figure S1: Example prior and posterior parameter values*

*The prior and posterior parameter values for a single scanner additive effect* $\gamma$*, corresponding to the additive scanner factor for the left entorhinal cortex from scanner 18. The interpretation of the exact* $\gamma$ *values across FB ComBat and EB ComBat differs slightly due to the standardization and scaling differences between the two methods. Thus, they are not comparable. However, they are expected to be directionally (i.e. positive or negative) similar.*

***Section S3: Scanner Strength Prediction***

|  | AUROC | |
| --- | --- | --- |
| Method | *Train* | *Test* |
| Unharmonized | 0.980 (0.003) | 0.949 (0.018) |
| EB ComBat | 0.868 (0.015) | 0.694 (0.040) |
| FB ComBat | 0.887 (0.009) | 0.735 (0.034) |

*Table S2: Scanner Strength Prediction Results*

*Scanner strength prediction performance using various harmonization methods and unharmonized data. Lower accuracy in the task indicates better harmonization performance due to more effective removal of non-biological (scanner magnetic field strength) information. EB ComBat performs best on this task (lowest test AUROC).*

***Section S4: Regional Uncertainty***

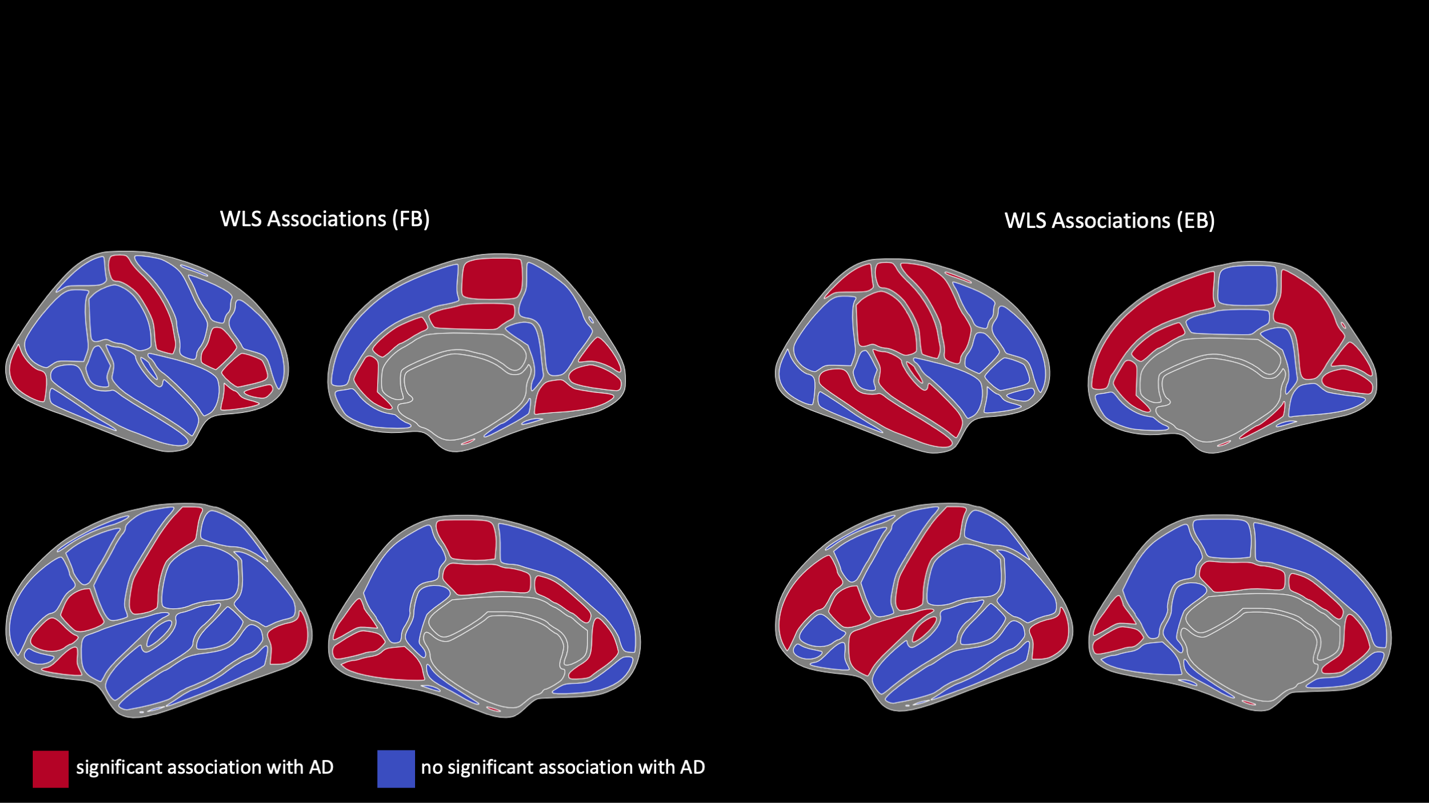

*Supplementary Figure S2: Cortical Thickness Associations with AD (FB vs. EB)*

*Cortical thickness regions associated with Alzheimer’s disease using Weighted Least Squares (WLS) weighted by the reciprocal of posterior measurement variance from FB ComBat (left) and EB ComBat (right).*

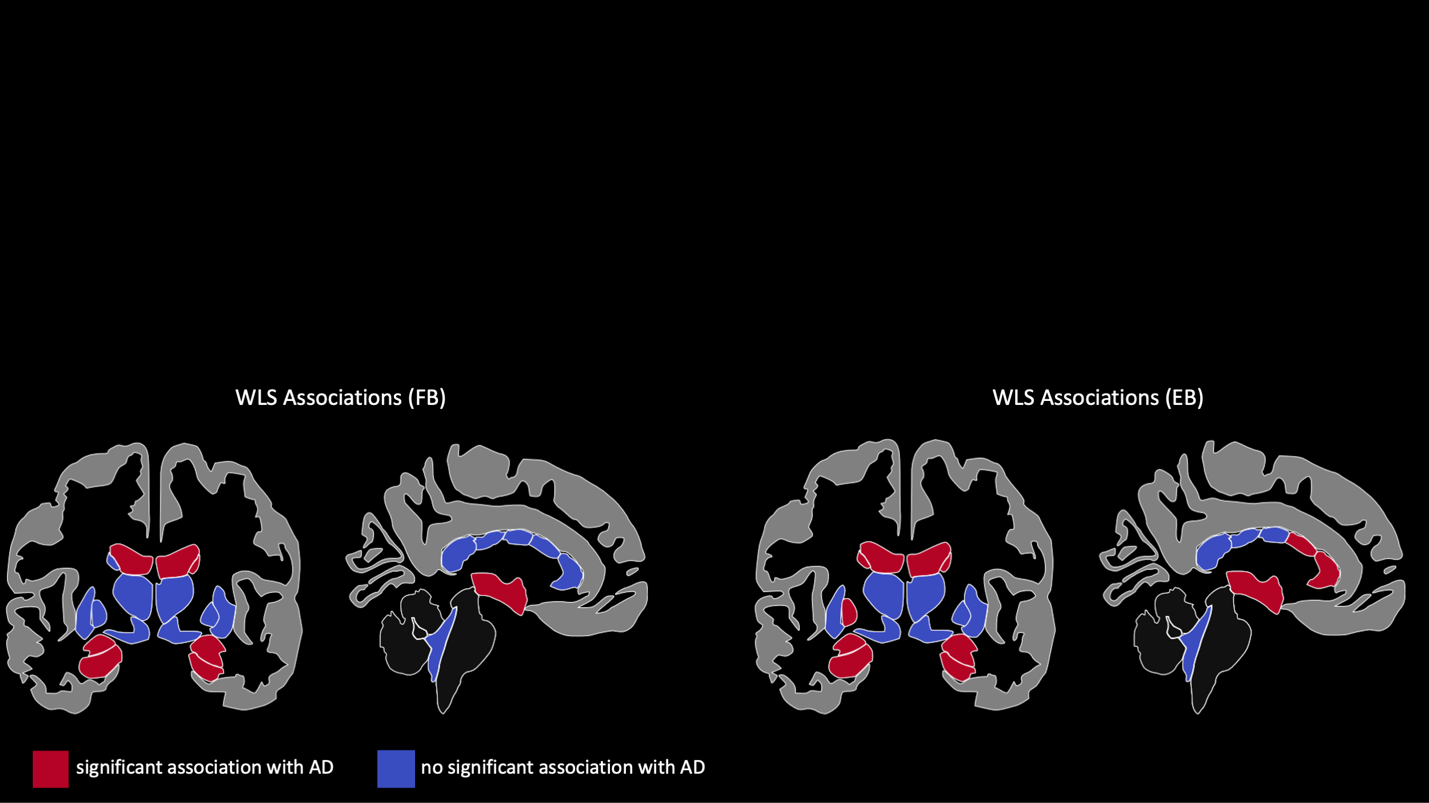

*Supplementary Figure S3: Subcortical Volume Associations with AD (FB vs. EB)*

*Subcortical volume regions associated with Alzheimer’s disease using Weighted Least Squares (WLS) weighted by the reciprocal of posterior measurement variance from FB ComBat (left) and EB ComBat (right).*

***Section S5: Simulation Study***

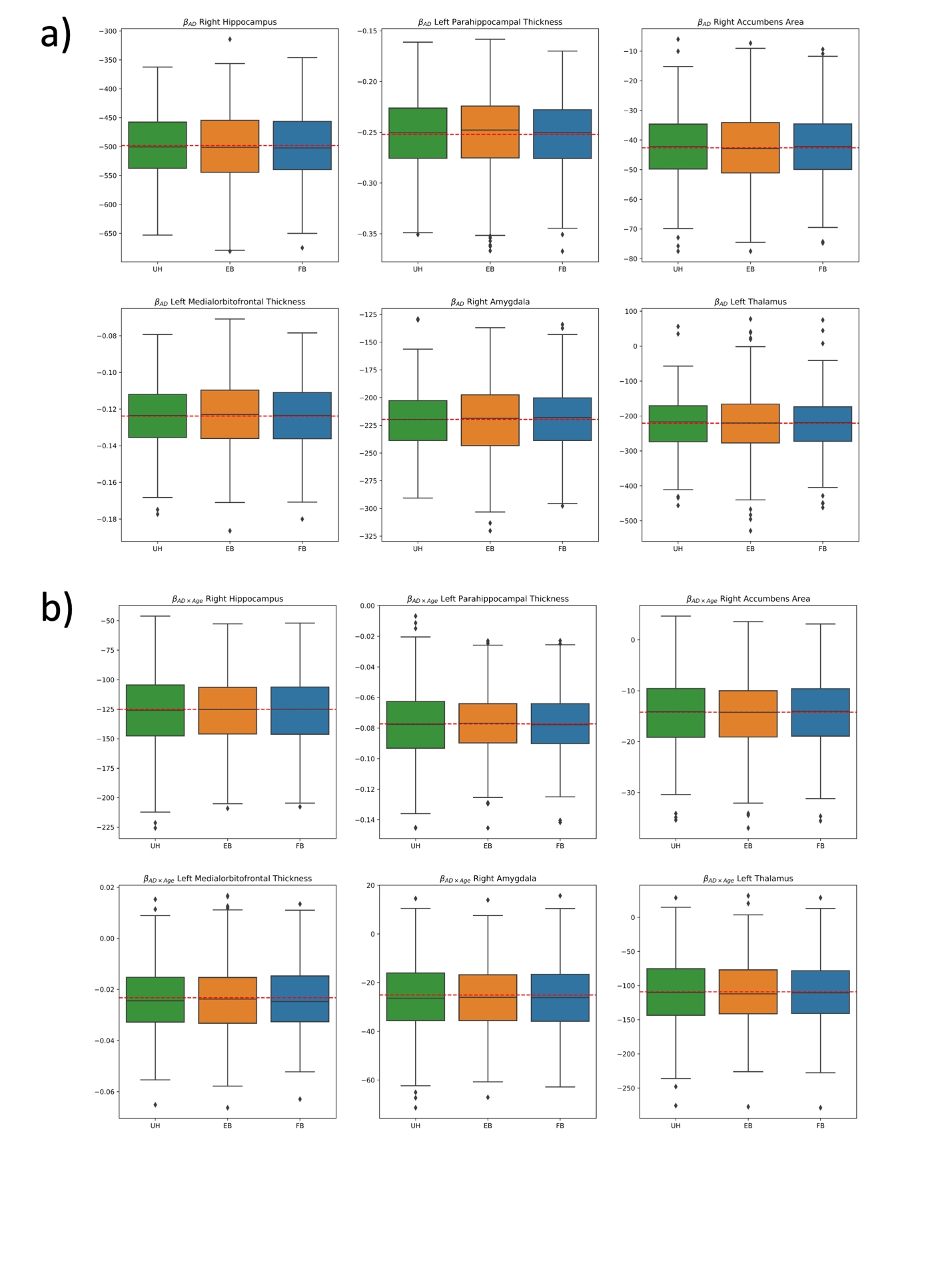

*Figure S4: Simulation Study Disease Effect Feature Coefficient Values*

*Simulation Data Coefficient Values for a) AD and b) AD x Age effect of the six true effect features. No significant difference in absolute error between different harmonization methods was found for any feature. The red dashed line indicates the true effect size.*

***Section S6: Sensitivity Analysis***

| **Experiment Name** | **Description** | **Changed Prior(s)** |
| --- | --- | --- |
| *Strong_Pooling* | Strong prior on $\tau_{i}$ towards 0 (i.e. strong pooling prior) | $\tau_{i} \sim IG(2,0.1)$ |
| *Strong_Pooling+* | Stronger prior on $\tau_{i}$ towards 0 (i.e. strong pooling prior) | $\tau_{i} \sim IG(2,0.01)$ |
| *Strong_No_Pooling* | Strong positive prior on $\tau_{i}$ (i.e. strong anti-pooling prior) | $\tau_{i} \sim G(4,2)$ |
| *Strong_No_Pooling+* | Strong positive prior on $\tau_{i}$ (i.e. strong anti-pooling prior) | $\tau_{i} \sim G(12,6)$ |
| *Normal_Mu* | Normal Prior on $\mu_{i}$ instead of Cauchy | $\mu_{i} \sim N(0,1)$ |
| *Strong_Priors* | *Priors are similar to baseline except much narrower. This simulates over-confident priors.* | $\mu_{i} \sim C\left( 0,0.01 \right)$  $\tau_{i} \sim IG\left( 11,5 \right)$  $\lambda_{i} \sim C\left( 1000,1000 \right)$  $\rho_{i} \sim IG\left( 26,25 \right)$  $\alpha_{v} \sim N\left( 0,0.01 \right)$  $\beta_{v} \sim N\left( 0,1 \right)$ |

*Table S3: Summary of Sensitivity Analysis Experiments. The first four explore whether a prior bias towards or against scanner pooling have any effect on harmonization performance. Normal_Mu examines the effect of Cauchy prior on the scanner additive parameter* $\mu_{i}$*. Strong_Priors replaces the baseline with stronger priors while keeping the prior mean (or mode for Cauchy distributions) constant.*

*Figure S5: Scanner Effects across Field Strength and Manufacturer in Sensitivity Analysis*

*Vertical lines show the distribution of normalized thickness and volume features for a single image; the region within one standard deviation of the image-specific mean is shaded darker. Horizontal lines show the means and 1 standard deviation intervals of normalized feature means for each scanner strength/manufacturer combination. Covariate effects are regressed out before plotting. Distribution distances across field strength and manufacturer are partially but not completely removed by EB ComBat, baseline FB ComBat, and all iterations of FB ComBat with modifications to priors. See Table S2 for description of prior specifications.*

*Figure S6: Linear Discriminant Analysis (LDA) with Respect to Field Strength and Manufacturer in Sensitivity Analysis*

*The first three LDA components of unharmonized and harmonized datasets (using the combination scanner strength and manufacturer as the target variable) are shown. EB ComBat and all variations of FB ComBat remove most scanner-related variation in LDA components. However, some scanner-related variation is still visible in all the FB ComBat specifications. Covariates were regressed out from features used for LDA, and these features were z-score normalized.*

| ***Harmonization Method*** | ***Signifigant (p<.05) Additive Effects (out of 122 features)*** | ***Significant (p<.05) Multiplicative Effects (out of 122 features)*** |
| --- | --- | --- |
| *Unharmonized* | *121* | *122* |
| *EB ComBat* | *0* | *1* |
| *FB ComBat* | *0* | *6* |
| *FB ComBat Normal_Mu* | *0* | *6* |
| *FB ComBat Strong_Priors* | *0* | *16* |
| *FB ComBat Strong_Pooling* | *0* | *6* |
| *FB ComBat Strong_Pooling+* | *0* | *6* |
| *FB ComBat Strong_No_Pooling* | *5* | *6* |
| *FB ComBat Strong_No_Pooling+* | *6* | *6* |

*Table S3: Significant additive and multiplicative scanner effects in sensitivity analysis*

*We calculate Kenward-Roger and Fligner-Killeen tests for unharmonized and harmonized datasets, including FB ComBat data from sensitivity analysis studies (See Table S1 for sensitivity study prior configurations).*
